# AUTS2 regulation of synapses for proper synaptic inputs and social communication

**DOI:** 10.1101/871012

**Authors:** Kei Hori, Kunihiko Yamashiro, Taku Nagai, Wei Shan, Saki F. Egusa, Kazumi Shimaoka, Yasuhiro Go, Shoji Tatsumoto, Mitsuyo Yamada, Reika Shiraishi, Kouta Kanno, Satoshi Miyashita, Asami Sakamoto, Manabu Abe, Kenji Sakimura, Masaki Sone, Kazuhiro Sohya, Hiroshi Kunugi, Kiyofumi Yamada, Mikio Hoshino

## Abstract

Impairments in synapse development are thought to cause numerous psychiatric disorders. *Autism susceptibility candidate 2* (*AUTS2*) gene has been associated with various psychiatric disorders, such as autism and intellectual disabilities. Although roles for AUTS2 in neuronal migration and neuritogenesis have been reported, its involvement in synapse regulation remains unclear. In this study, we found that excitatory synapses were specifically increased in the *Auts2*-deficient primary cultured neurons as well as *Auts2* mutant forebrains. Electrophysiological recordings and immunostaining showed increases in excitatory synaptic inputs as well as c-fos expression in *Auts2* mutant brains, suggesting that an altered balance of excitatory and inhibitory inputs enhances brain excitability. *Auts2* mutant mice exhibited autistic-like behaviors including impairments in social interaction and altered vocal communication. Together, these findings suggest that AUTS2 regulates excitatory synapse number to coordinate E/I balance in the brain, whose impairment may underlie the pathology of psychiatric disorders in individuals with *AUTS2* mutations.

## [Introduction]

Synapses form the basis for the neuronal network and brain function. Development of synapses, synaptogenesis, is precisely regulated by genetic programs as well as synaptic activities. Even after establishment of the fundamental brain structures, synapses are dynamically formed and eliminated in response to neuro-environmental stimuli (Holtmaat and Svoboda, 2009). However, maintenance of the number of synapses within a certain range, comprising the synapse homeostasis, assures neuronal homeostasis (Davis, 2013; Tien and Kerschensteiner, 2018; Wefelmeyer et al., 2016). It has been proposed that failure of either synapse or neuronal homeostasis results in various neuropsychiatric disorders (Bourgeron, 2009; Ramocki and Zoghbi, 2008). Consistent with this, postmortem pathological studies have revealed that aberrant regulation of dendritic spine number as well as structural abnormalities of spines were observed in patients with numerous psychiatric disorders such as autism spectrum disorders (ASDs), schizophrenia and neurodegenerative diseases (Hutsler and Zhang, 2010; Penzes et al., 2011; Tang et al., 2014). Thus, appropriate regulation of synaptogenesis as well as synapse homeostasis is critical for normal healthy brain function, however, its molecular machinery remains elusive.

*Autism susceptibility candidate 2* (*AUTS2*) (also termed “activator of transcription and developmental regulator”) located on human chromosome 7q11.22 has been initially identified as a possible ASD risk gene in a study that reported a *de novo* balanced translocation in monozygotic twin patients with ASDs (Sultana et al., 2002). Thereafter, structural variants that disrupt the *AUTS2* locus have been identified in the patients with not only autism, but also other neuropathological conditions including intellectual disabilities (IDs), schizophrenia, attention deficit hyperactivity disorder (ADHD), dyslexia and epilepsy as well as brain malformation and craniofacial abnormalities (Amarillo et al., 2014; Bakkaloglu et al., 2008; Ben-David et al., 2011; Beunders et al., 2013; Elia et al., 2010; Hori and Hoshino, 2017; Jolley et al., 2013; Kalscheuer et al., 2007; Oksenberg and Ahituv, 2013; Talkowski et al., 2012; Zhang et al., 2014). In addition, *AUTS2* has been recently implicated as a potential gene in human-specific evolution (Oksenberg and Ahituv, 2013; Oksenberg et al., 2013).

We previously reported that AUTS2 acts as an upstream regulator for Rho family small GTPases, Rac1 and Cdc42, in reorganizing actin cytoskeleton (Hori et al., 2014). AUTS2 activates Rac1 to induce lamellipodia, while downregulating CDC42 to suppress filopodia. In addition to these functions, Gao *et al* showed that AUTS2 binds to and neutralizes the transcriptional repressor activity of Polycomb group (PcG) protein complex 1 (PRC1) and activates some gene transcription by recruiting the histone acetyltransferase P300 into the complex (Gao et al., 2014).

In the developing mouse brain, *Auts2* expression starts from early embryonic stages in multiple regions of the central nervous system, but particularly strong prenatal expression is observed in the regions associated with higher brain functions including neocortex, hippocampus and cerebellum (Bedogni et al., 2010). We previously demonstrated that the AUTS2-Rac1 signaling pathway is required for neuronal migration and subsequent neurite formation in the developing cerebral cortex (Hori et al., 2014). However, even at postnatal and adult stages, AUTS2 expression is maintained in various types of neurons (Bedogni et al., 2010). Although this late-stage expression raised the possibility that AUTS2 may also be involved in later neurodevelopmental processes, such as synaptogenesis and synaptic homeostasis, its involvement in synapse regulation remains unknown.

In human patients, *AUTS2* mutations are associated with a variety of psychiatric diseases, such as ASD, schizophrenia, depression, intellectual disabilities and language disability. Although the underlying pathways to evoke this wide range of disorders have not been clarified, one possible mechanism is that different types of gene disruption may cause distinct types of disorders. *AUTS2* is a very large gene with multiple exons and many types of gene mutations, such as deletion, duplication, single nucleotide change, and chromosomal translocation have been reported in humans (Hori and Hoshino, 2017; Oksenberg and Ahituv, 2013).

In this study, we show that AUTS2 coordinates excitation/inhibition balance, by restricting the number of excitatory synapses during development as well as at post-developmental stages. Targeted disruption of *Auts2* resulted in excessive numbers of excitatory synapses without affecting inhibitory ones. Consistent with this, electrophysiological analyses showed that excitatory but not inhibitory inputs increased in the mutant hippocampal neurons where strong c-Fos signals were detected, suggesting impairment in the excitatory and inhibitory coordination in that region. We performed behavioral analyses on heterozygotes for another *Auts2* allele, *Auts2^del8^*, whose AUTS2 expression profile is distinct from that of *Auts2^neo^* (Supplementary Table 1) (Hori et al., 2015). and found that *Auts2^neo/+^* and *Auts2^del8/+^* exhibited overlapping but distinct behavioral abnormalities including social interaction, resembling a wide range of psychiatric symptoms in individuals with *AUTS2* mutations. Thus, our data suggests that AUTS2 regulates synapse homeostasis by restricting the number of excitatory synapses without affecting inhibitory ones and that loss of AUTS2 function leads to impaired excitatory and inhibitory coordination that may underlie the pathogenesis of some psychiatric illnesses.

## [Results]

### *Auts2* restricts the number of excitatory synapses *in vitro*

To investigate the involvement of AUTS2 in synapse formation, we utilized primary cultured hippocampal neurons from homozygous *Auts2-floxed* (*Auts2^flox/flox^*) embryos. Most excitatory synapses in mammalian brain are formed on dendritic spines (Bhatt et al., 2009). We confirmed that at 21 days *in vitro* (DIV 21), most PSD-95 (excitatory postsynapse marker) signals were observed on the spine heads (Figure 1A).

**Figure 1.**
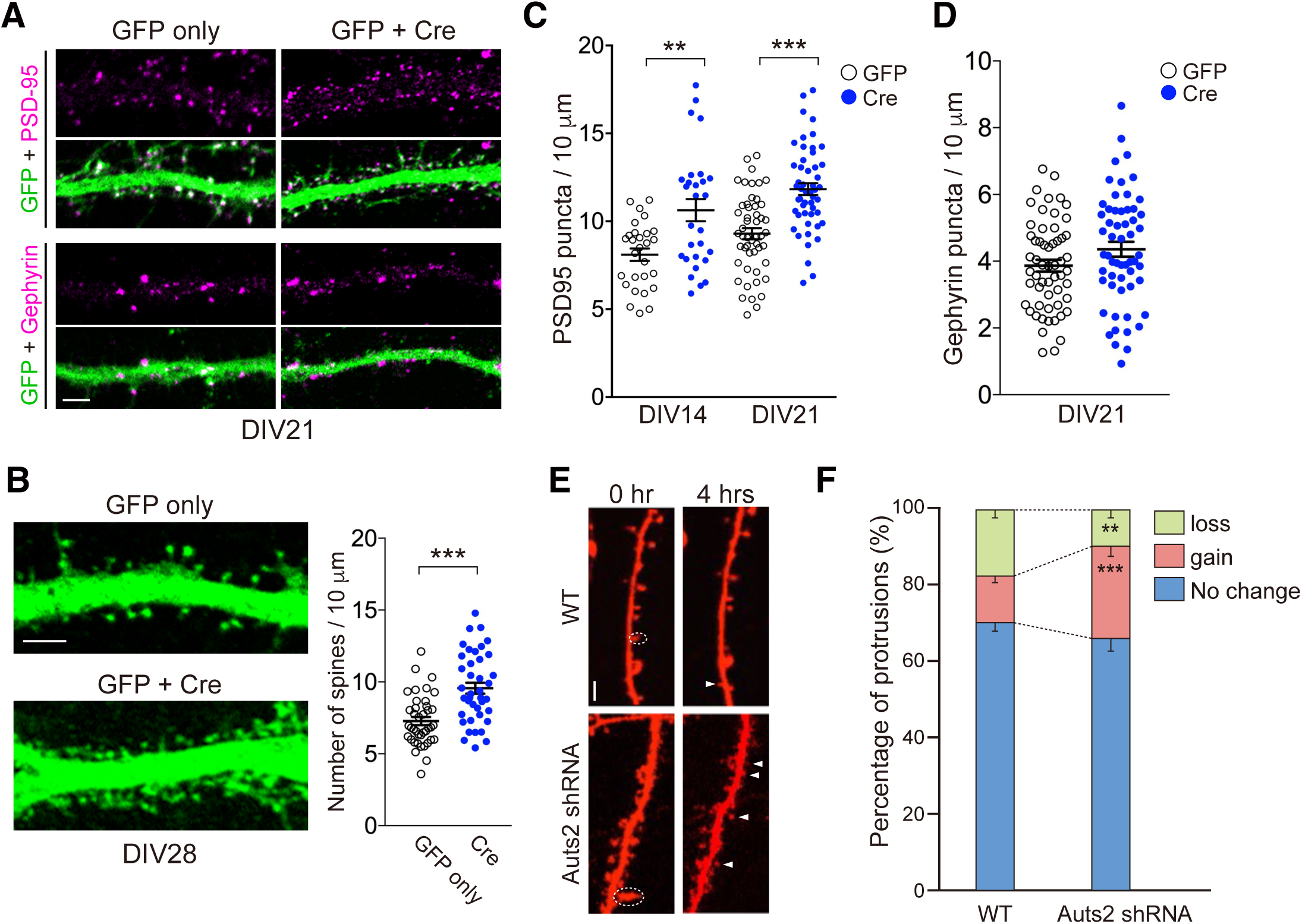
Loss of *Auts2* induces excessive excitatory synapse formation. (A) Primary hippocampal neurons derived from *Auts2^flox/flox^* homozygotes at DIV21 were immunolabeled with anti-GFP (green) and anti-PSD-95 or Gephyrin (magenta). Neurons were co-electroporated with control or Cre plus GFP expression vector. (B) Dendritic spines were increased in *Auts2* KO neurons (GFP + Cre) at DIV28. The graph shows the density of dendritic spines in the GFP-positive neurons. (n=40 dendrites from 20 neurons). (C and D) The number of PSD-95 (C) and gephyrin (D) postsynaptic puncta colocalized with GFP was measured at DIV14 and 21 (DIV14, n=28 dendrites; DIV21, n=51 dendrites of 15-22 neurons in B and n=57 dendrites of 19 neurons in (C). (E) WT mouse hippocampal neurons at DIV16 transfected with mRFP only (WT) or mRFP plus Auts2 shRNA vector were imaged at the beginning (0h) and 4 hrs after the analysis. (Dashed white circle, spine eliminated; white arrowheads, spines formed). (F) Gain and loss of dendritic protrusions (including spines and filopodia) in WT and *Auts2* knockdown neurons were analyzed during a 6 hr time window at DIV16-17. (WT, n=7 neurons; *Auts2* shRNA, n=10 neurons). Data are presented as mean ± SEM. Quantifications in B-D and F represent data from three independent experiments. \*\**P* < 0.01, \*\*\**P* < 0.001, (B-D) unpaired t-test, (F) Mann-Whitney U test. Scale bars represent 10 μm.

Deletion of *Auts2* was carried out by co-introducing GFP with the Cre recombinase expression vector into the *Auts2^flox/flox^* primary hippocampal neurons. Immunostaining revealed that the *Auts2*-deficient neurons (*Auts2^del8/del8^* neurons) exhibited a significant increase in the density of dendritic spines compared with the control neurons at DIV28 (Figure 1B). Consistent with the increased dendritic spines, *Auts2*-deficient neurons harbored a larger number of PSD-95 puncta than the control at DIV 21 (Figure 1A and C). The larger numbers of PSD-95 puncta were already evident at an early culture stage (DIV 14) in the mutant neurons (Figure 1C). Interestingly, the number of an inhibitory post synapse marker, Gephyrin-positive puncta was not different between the control and *Auts2*-deficient neurons (Figures 1A and D). These findings suggest that *Auts2* in postsynaptic cells restricts excessive excitatory synapse formation without affecting inhibitory synapses.

We further observed the development of dendritic spines at different stages in culture (Supplementary Figure 1A). In control neurons, filopodia were predominantly formed during the first week of culture but gradually decreased from 2 to 4 weeks, with increasing spine formation during the same period. During the first week of culture, *Auts2* mutant neurons had a similar number of protrusions including filopodia and spines as control neurons (Supplementary Figure 1A). At later stages, however, larger numbers of dendritic spines as well as filopodia were continuously formed in the *Auts2* mutant neurons compared to the control neurons (Supplementary Figure 1A). We measured the length of spines as well as the width of the spine heads as indices of spine maturity. There were no significant differences in either index, suggesting that loss of *Auts2* does not influence the maturation of dendritic spines (Supplementary Figure 1B).

Next, we introduced the expression vectors for AUTS2 isoforms or possible AUTS2 downstream factors into the *Auts2*-knockdown neurons (Hori et al., 2014). We first confirmed that knockdown of *Auts2* well recapitulated aberrant spine formation as observed in *Auts2* KO neurons (Supplementary Figure 1C). This abnormality was restored by co-expression of the shRNA-resistant full-length AUTS2 (FL-AUTS2^R^), indicating that excess spine formation is the result of specific knockdown of *Auts2* (Figure 1C). We have previously demonstrated that a cytoplasmic AUTS2-Rac1 signaling pathway is required for neuronal migration in the developing cerebral cortex (Hori et al., 2014). In that study, defective cortical neuronal migration in *Auts2* KO mice was shown to be rescued by introduction of either NES (nuclear export sequence)-tagged FL-AUTS2^R^ (NES-FL-AUTS2^R^) or wild type Rac1 (Rac1-WT). Overexpression of these proteins, however, failed to rescue the aberrant spine formation evoked by *Auts2* knockdown (Supplementary Figure 1C). In contrast, introduction of NLS (nuclear localization signal)-tagged FL-AUTS2^R^ (NLS-FL-AUTS2^R^) was able to rescue the spine number to levels comparable to that of control neurons, while the C-terminal AUTS2 short isoform (S-AUTS2-var.1) was not able to rescue the phenotype (Supplementary Figure 1C and 2B). These results indicate that nuclear FL-AUTS2 is involved in the control of spine number.

Next, we performed live imaging to observe the dynamics of dendritic protrusions including spines and filopodia at DIV 16-17. During a 6 hr time window, neurons expressing the *Auts2* shRNA exhibited a higher rate of protrusion gain and a lower rate of protrusion loss compared with WT neurons (Figure 1E and F). Altogether, these *in vitro* experiments suggest that AUTS2 restricts the number of excitatory synapses by suppressing spine formation and/or by eliminating pre-existing spines, while not affecting inhibitory neurons.

### Loss of *Auts2* results in excessive dendritic spines *in vivo*

We generated forebrain-specific *Auts2* conditional KO mice by crossing *Auts2-floxed* mice with *Emx1^Cre^* mice (Iwasato et al., 2000) (Supplementary Figures 2A and Supplementary Table 1) and examined brain tissues by Golgi staining to visualize dendrite morphology. Immunoblotting confirmed that expression of FL-AUTS2 protein in the mutant cerebral cortex was successfully eliminated (arrow in Supplementary Figure 2C). We found that dendritic spines were increased in the dendrites of layer II/III pyramidal neurons in the medial prefrontal cortex (mPFC) at P35 in *Emx1^Cre/+^;Auts2^flox/flox^* homozygous mutant brains compared with the *Auts2^flox/flox^* controls (Figures 2A and B). Significant differences were not limited to mPFC neurons. For example, increased spines were also observed in apical dendrites of hippocampal CA1 pyramidal neurons and dendrites of the upper-layer neurons of the auditory cortex (Figure 2B and Supplementary Figure 2D). These findings suggest that AUTS2 restricts the number of dendritic spines *in vivo*. Interestingly, a similar phenotype was observed in *Emx1^Cre/+^;Auts2^flox/+^* heterozygous mutants (Figure 2B).

**Figure 2.**
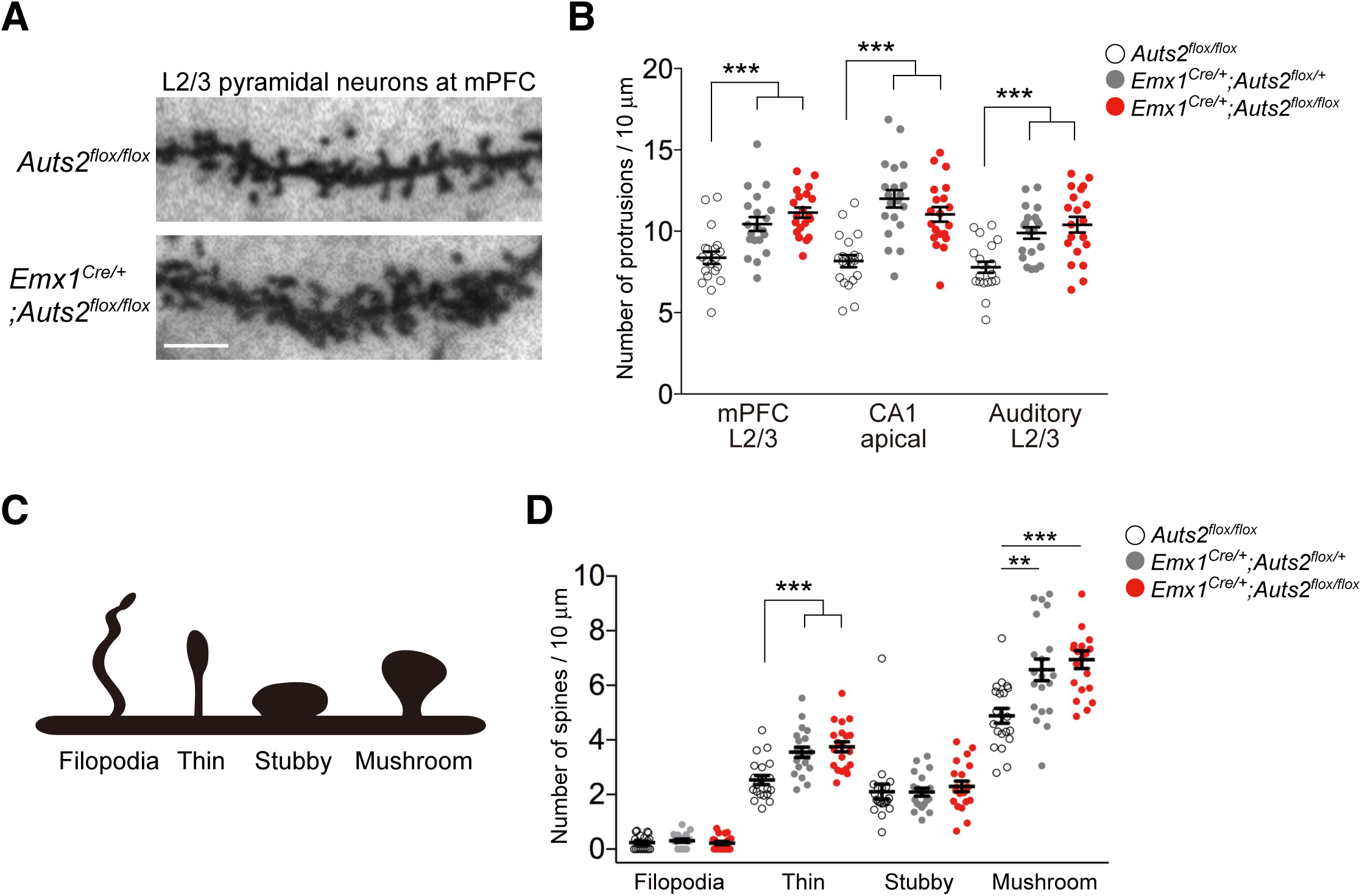
Loss of *Auts2* abnormally increases dendritic spine formation *in vivo*. (A) Representative images of the dendritic spines from Golgi-stained upper-layer pyramidal neurons in the mPFC of control (*Auts2^flox/flox^*, upper panel) and *Emx1^Cre/+^;Auts2^flox/flox^* homozygous mutant mouse brains (lower panel) at P35. (B) Summary graph of the spine density on the neurons at indicated areas in the control (*Auts2^flox/flox^*), heterozygous (*Emx1^Cre/+^;Auts2^flox/+^*) and homozygous (*Emx1^Cre/+^;Auts2^flox/flox^*) mutant mouse brains. (n=20 dendrites from n=3 animals). (C) Morphological classification of dendritic spines and filopodia. (D) The density of each category of spines in the upper-layer neurons in the mPFC was measured in the control (*Auts2^flox/flox^*), heterozygous (*Emx1^Cre/+^;Auts2^flox/+^*) and homozygous (*Emx1^Cre/+^;Auts2^flox/flox^*) mutant mouse brains. (n=20 dendrites from n=3 animals). Data are presented as mean ± SEM. \*\**P* < 0.01, \*\**P <* 0.001, one-way ANOVA with Dunnett’s post hoc test. Scale bar, 10 μm.

We categorized spines into four morphological types (Figure 2C) and found that both mature “mushroom” spines and immature “thin” spines were increased to a similar extent in dendrites of *Emx1^Cre/+^;Auts2^flox/+^* heterozygous and *Emx1^Cre/+^;Auts2^flox/flox^* homozygous neurons (Figure 2D). This indicates that AUTS2 does not affect the maturity of spines, as was also observed in our *ex vivo* data (Supplementary Figure 1B). These observations suggest that AUTS2 restricts the number of excitatory synapses and that loss of one allele is sufficient to result in excessive excitatory synapses.

### *Auts2* deficiency causes aberrant excitatory neurotransmission

Next, we investigated the effect of *Auts2* inactivation on synaptic transmission properties. To address this, we performed whole-cell patch clamp recording of spontaneous miniature excitatory and inhibitory postsynaptic currents (mEPSCs and mIPSCs, respectively) on CA1 pyramidal neurons in acute hippocampal slices from P33-44 mouse brains. In the *Emx1^Cre/+^;Auts2^flox/flox^* homozygous brains, the mEPSCs were increased in frequency but not in amplitude compared with the control (*Auts2^flox/flox^*) mice (Figures 3A and C). On the other hand, no difference was observed in the mIPSCs with regard to either amplitude or frequency between the control and *Emx1^Cre/+^;Auts2^flox/flox^* mutants (Figures 3B and D). These results, together with the increase in spine density and PSD-95 puncta, suggest that the increased frequency of spontaneous mEPSCs in *Auts2* mutant pyramidal neurons might reflect excessive input from increased excitatory synaptic contacts.

**Figure 3.**
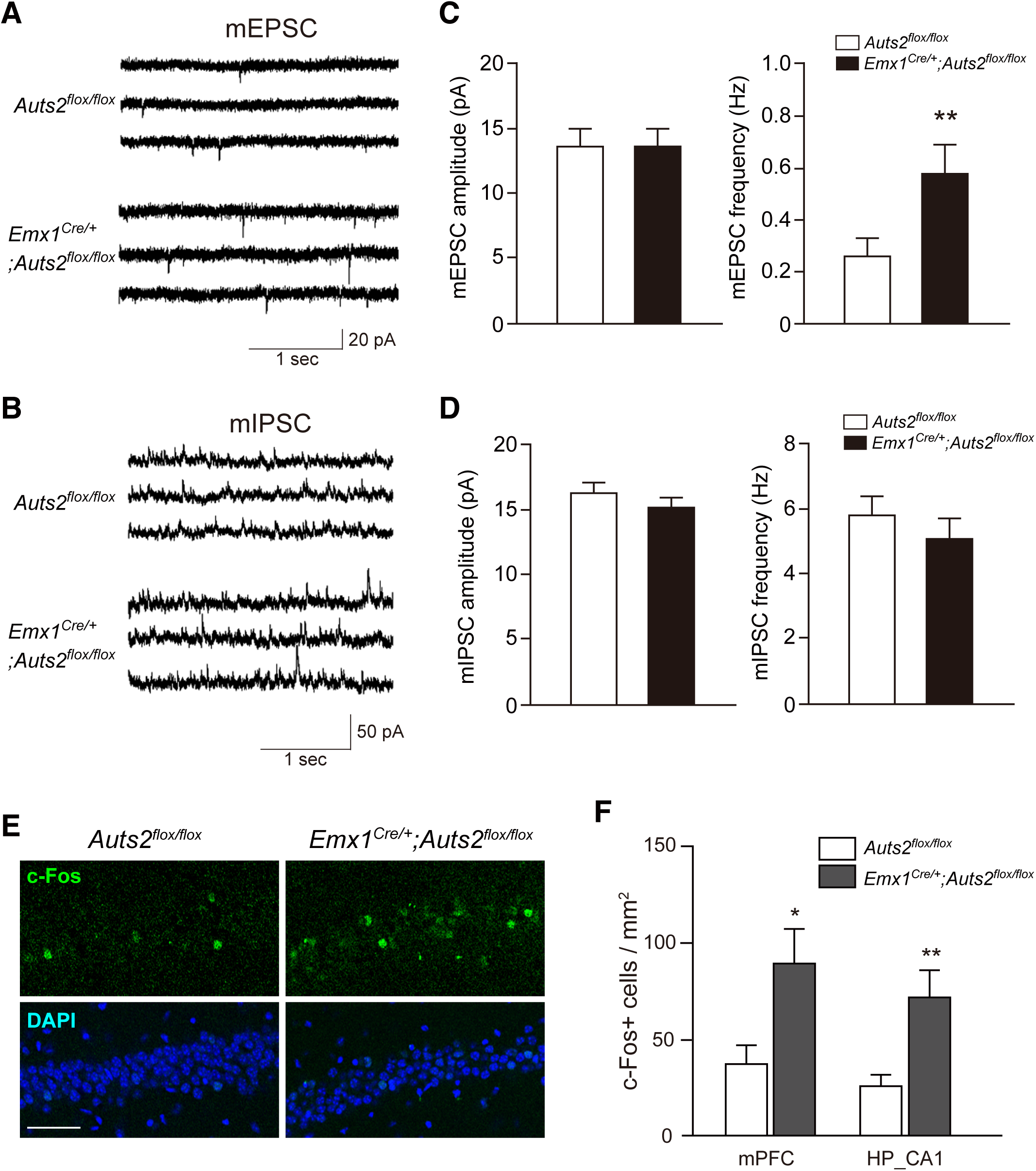
*Auts2* mutant mice display altered synaptic properties and increased c-Fos expression. (A, B) Representative traces of mEPSCs (A) and mIPSCs (B) from slice recordings of CA1 pyramidal neurons from control (*Auts2^flox/flox^*) and *Emx1^Cre/+^;Auts2^flox/flox^* homozygous mutant mice at P35. (C, D) *Emx1^Cre/+^;Auts2^flox/flox^* mice exhibit increased mEPSC (C) but not mIPSC (D) frequencies without change in amplitude. n=18-19 neurons from N=6-8 mice per genotype. (E) Representative images of c-Fos expression in the hippocampal CA1 areas of homozygous *Emx1^Cre/+^;Auts2^flox/flox^* homozygous mutant mice and *Auts2^flox/flox^* control littermates. (F) Summary graphs of c-Fos-expressing cells in the indicated areas. 8-12 tissue sections from N=3 brains were analyzed. Data are presented as Mean ± SEM. \**P* < 0.05, \*\**P* < 0.01, Mann-Whitney U test. Scale bar, 50 μm.

Furthermore, we examined the expression of the immediate-early gene product, c-Fos, as a marker of neuronal activity in the brain (Sagar et al., 1988). Compared with the control (*Auts2^flox/flox^*) mice, a larger number of pyramidal neurons with strong c-Fos immunoreactivity were observed in the mPFC and hippocampal CA1 in *Emx1^Cre/+^;Auts2^flox/flox^* homozygous mutants (Figures 3E and F). This suggests that the disturbed balance between excitatory and inhibitory synaptic inputs in local neural circuits results in increased excitability in the *Auts2* mutant brains.

### *Auts2* prevents excessive spine formation even after developmental stages

Although our *ex vivo* and *in vivo* analyses suggest that AUTS2 regulates excitatory synapse formation, it is unclear whether AUTS2 possesses such a function after establishment of brain structures. To assess this issue, we crossed *Auts2-floxed* mice with *CaMKIIa-CreER^T2^* mice to generate *CaMKIIa-CreER^T2^;Auts2^flox/flox^* homozygous mutant mice, in which the exon 8 of *Auts2* can be ablated in the forebrain projection neurons by administration of tamoxifen (Erdmann et al., 2007) (Supplementary Figure 3A and Supplementary Table 1). Tang et al. previously demonstrated that the *CaMKIIa*-promoter is active in forebrain neurons from postnatal week 3 to adulthood (Tang et al., 2014). When tamoxifen was administered during P21-25 to *CaMKIIa-CreER^T2^;Auts2^flox/flox^* mutant mice and their control littermates (*Auts2^flox/flox^*), genomic recombination was detected in the mPFC and hippocampus but not in the cerebellum of *CaMKIIa-CreER^T2^;Auts2^flox/flox^* mice (Supplementary Figure 3B), indicating that this protocol efficiently induces the forebrain-specific Cre-mediated recombination. Induction of recombination was also confirmed by using *Rosa26R^YFP^*, a reporter allele to detect Cre-dependent recombination (Supplementary Figure 3C). Quantitative RT-PCR revealed that *Auts2* mRNA levels dramatically decreased in the mPFC and hippocampus but not in the cerebellum of the tamoxifen-treated *CaMKIIa-CreER^T2^;Auts2^flox/flox^* mice (Supplementary Figure 3D).

Three weeks after tamoxifen administration to *CaMKIIa-CreER^T2^;Auts2^flox/flox^* homozygous mutants and *Auts2^flox/flox^* control mice (Figure 4A), *CaMKIIa-CreER^T2^;Auts2^flox/flox^* mice displayed an increase in the densities of spines on the dendrites of both cortical and hippocampal pyramidal neurons (Figures 4B and C). Similar to the *Emx1^Cre/+^;Auts2^flox/flox^* mutant mice, those increased spines consisted of mushroom and stubby-type mature spines as well as immature thin spines (Figure 4D). These findings suggest that AUTS2 is required for the dendritic spine number restriction even at post-developmental stages, which may contribute to the regulation of synaptic homeostasis.

**Figure 4.**
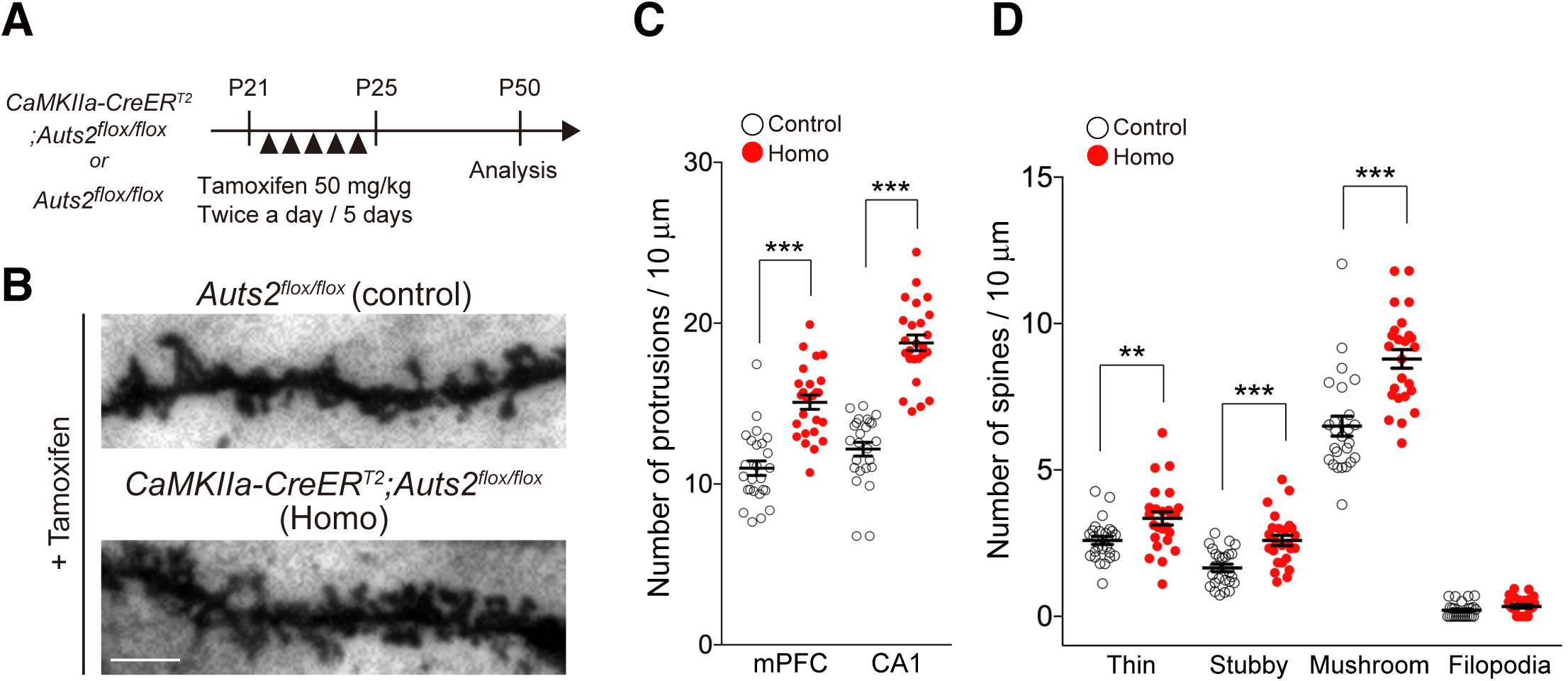
Conditional deletion of *Auts2* in postnatal forebrain leads to excessive spine formation. (A) Scheme illustrating the tamoxifen-inducible deletion of *Auts2* in postnatal forebrain. Tamoxifen was administered to *CaMKIIa-CreER^T2^;Auts2^flox/flox^* homozygotes and their control *Auts2^flox/flox^* littermate mice during P21-25 and analysis was performed at P50. (B) Representative images of the dendritic spines from Golgi-stained upper-layer pyramidal neurons at mPFC of the tamoxifen-treated control (*Auts2^flox/flox^*, upper panel) and *Auts2* homozygous mutant mouse brains (*CaMKIIa-CreER^T2^;Auts2^flox/flox^*, lower panel) at P50. (C) The pyramidal neurons in the mPFC as well as hippocampal CA1 area from mice postnatally lacking *CaMKIIa-CreER^T2^;Auts2^flox/flox^* (Homo) exhibited increase of dendritic spines relative to the *Auts2^flox/flox^* littermates (control). (D) The density of each category of spines on the pyramidal neurons in the mPFC was measured in control (*Auts2^flox/flox^*) and homozygous *CaMKIIa-CreER^T2^;Auts2^flox/flox^* mutant mouse brains (n=25 dendrites from N=3 animals). Data are presented as mean ± SEM. \*\**P* < 0.01, \*\*\**P* < 0.001, unpaired t-test. Scale bar, 10 μm.

### Aberrant gene expression in *Auts2* mutant mice

The *ex vivo* rescue experiments in Supplementary Figure 1C showed that AUTS2 in the nucleus functions to restrict the spine number. A previous study clarified that nucleic AUTS2 works as a component of PRC1 to participate in gene transcription (Gao et al., 2014). These findings suggest that AUTS2 protein in nuclei restricts spine formation by regulating gene expression of relevant neural genes. Therefore, we examined global mRNA expression profiles for *Emx1^Cre/+^;Auts2^flox/flox^* homozygous brains and *Auts2^flox/flox^* control littermate brains. Through RNA-sequencing, we identified a total of 168 genes, whose expression levels were significantly altered in the mutant hippocampus, with 78 downregulated and 90 upregulated genes (Figure 5A-C). Interestingly, these differentially expressed genes included the genes encoded synaptic proteins or molecules involved in synaptic functions, such as *Reln*, *Mdga1*, *Camk2b*, *Cacna1c* and *C1ql*-family genes (Fink et al., 2003; Gangwar et al., 2017; Martinelli et al., 2016; Matsuda et al., 2016; Moosmang et al., 2005; Pettem et al., 2013; van Woerden et al., 2009; Wasser and Herz, 2017) (Figure 5B and C). Gene ontology (GO) analysis revealed that these altered genes were associated with multiple aspects of neurodevelopment including “nervous system development”, “cell differentiation”, “neuronal migration”, with particular enrichment of the terms for synapse development such as “dendritic spine morphogenesis”, “negative regulation of synapse assembly” and “regulation of cytosolic calcium ion concentration” (Figure 5D). These results suggest that nucleic AUTS2 regulates the expression of genes that are related to synapse formation/function, and some of which may be involved in spine number restriction. Aberrant expression of such synaptic genes may cause synaptic dysfunction in patients with *AUTS2* mutations.

**Figure 5.**
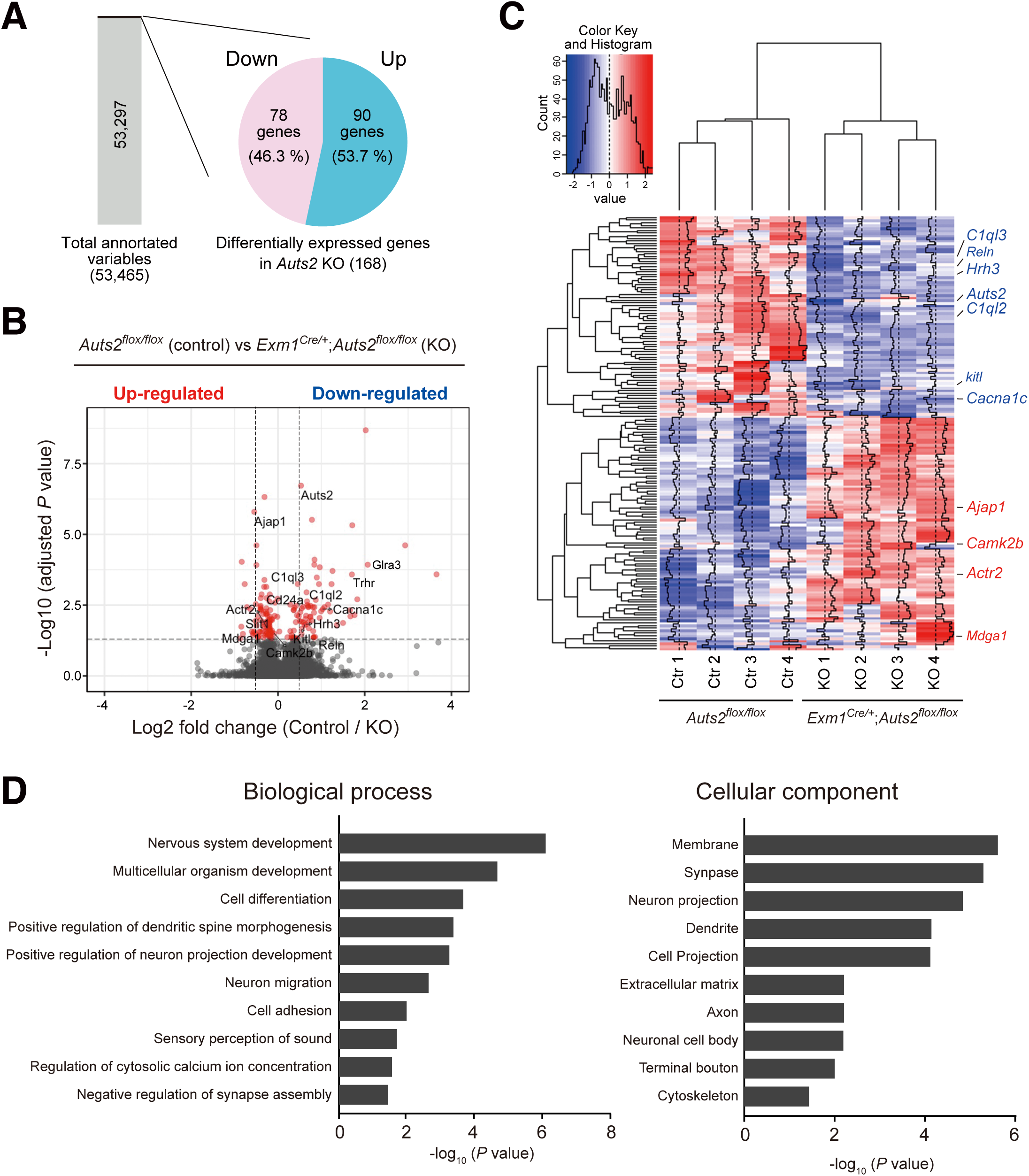
Transcriptome analysis of *Emx1^Cre/+^;Auts2^flox/flox^* mutant mice hippocampal brain tissues. Global gene expression analysis by RNA-sequencing reveals dysregulation of multiple genes associated with neurodevelopment. RNA samples from P14 hippocampus of *Emx1^Cre/+^;Auts2^flox/flox^* homozygous mutant mice and the *Auts2^flox/flox^* control littermates were used. **(A)** Rates in differentially expressed genes in *Emx1^Cre/+^;Auts2^flox/flox^* homozygous mutant hippocampal tissues compared to the *Auts2^flox/flox^* control littermates. (B) Volcano plot showing differential expression of all genes between *Auts2^flox/flox^* (control) and *Emx1^Cre/+^;Auts2^flox/flox^* homozygous mutants (KO). *P* value threshold of 0.05 for the false discovery rate (FDR), and of 0.5 for log_2_ fold change (log2FC) were indicated by horizontal and vertical dashed lines, respectively. **(C)** Clustered heatmap of transcriptome analysis in *Emx1^Cre/+^;Auts2^flox/flox^* homozygous mutants (KO) and the *Auts2^flox/flox^* control littermates (Ctr). Four biological samples as indicated were subjected to RNA-seq analysis. **(D)** Gene ontology (GO) analysis of the differentially expressed genes in *Auts2* mutant hippocampus.

### Behavioral abnormalities in *Auts2* mutant mice

All experimental mice including *Auts2^del8/+^* mutants (Figure 6 and Supplementary Figure 4), tamoxifen-treated *CaMKIIa-CreER^T2^;Auts2^flox/flox^* mice (Supplementary Figure 5) and *Auts2^flox/flox^* control littermates appeared grossly normal. All of them had normal fur and whiskers and showed no detectable motor disability. The body weight of *Auts2^del8/+^* mice was slightly decreased compared with WT littermates (body weight at 3 months of age; WT, 27.94 ± 0.54 g (n=16); *Auts2^del8/+^*, 20.50 ± 0.35 (n=16); data are mean ± SEM, Mann-Whitney *U* = 2.5, \*\*\**P* < 0.001).

**Figure 6.**
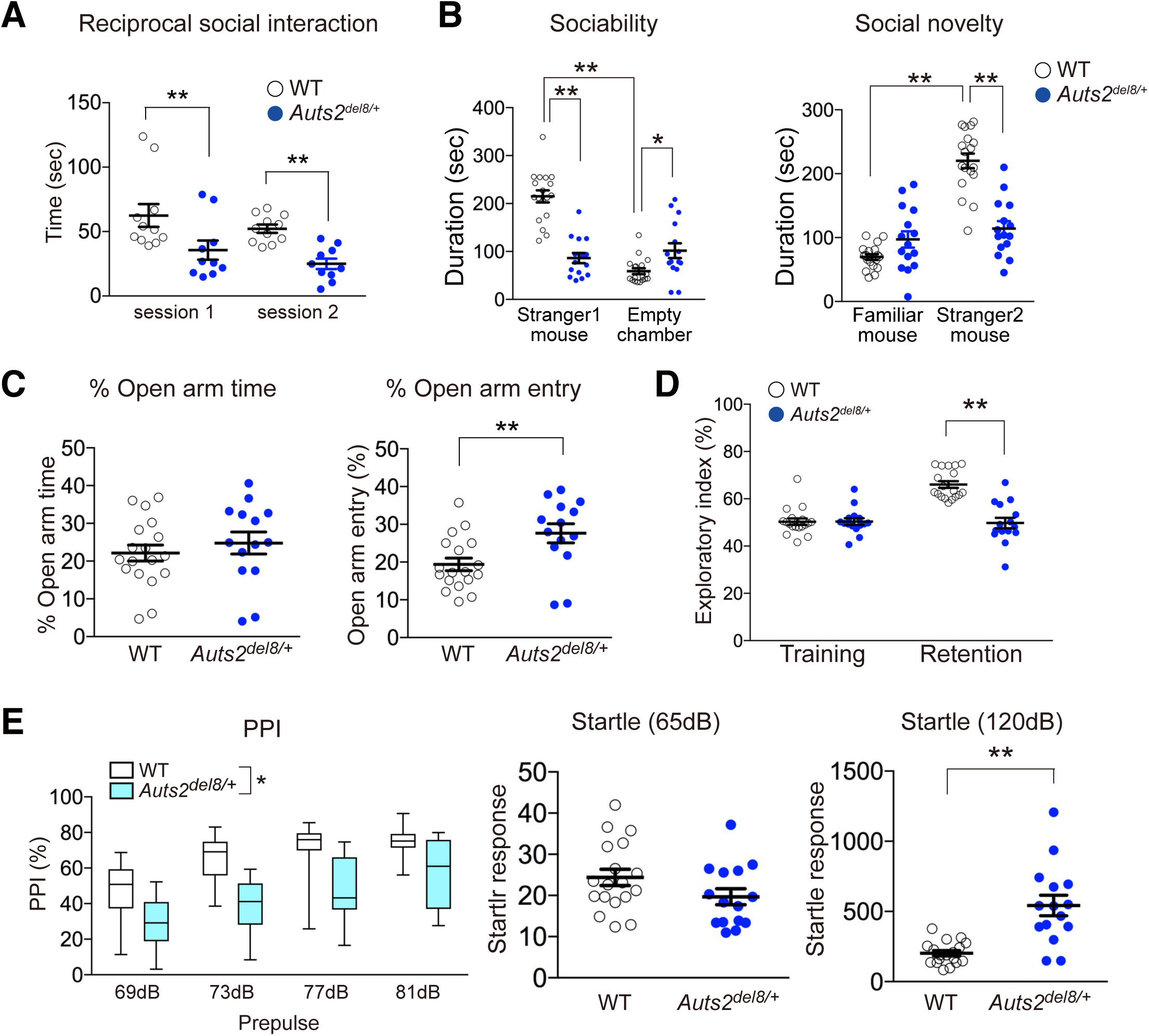
Behavioral abnormalities in *Auts2^del8/+^* mutant mice. (A) Reciprocal social interaction test. Social interaction between WT or *Auts2^del8/+^* mouse pairs during 5 min were measured (WT, n=11, *Auts2^del8/+^*, n=10). (B) Three-chamber social interaction test. Graphs show the amount of time spent in each chamber (WT, n=18, *Auts2^del8/+^*, n=15). (C) *Auts2 ^del8/+^* mice exhibit increased open arm entry relative to WT mice in elevated plus maze test (WT, n=18, *Auts2^del8/+^*, n=14). (D) *Auts2 ^del8/+^* mice display deficits in novel object recognition. Graphs show the exploratory preference in training and retention sessions (WT, n=18, *Auts2 ^del8/+^*, n=15). (E) Prepulse inhibition (PPI) (%) at four different prepulse intensities in PPI test (left graph) and acoustic startle response (middle and right graphs) as measured in trials without a prepulse. *Auts2 ^del8/+^* mice display decrease of the percentage of PPI as well as a higher acoustic startle response at 120 dB pulse relative to those in WT mice (WT, n=18, *Auts2^del8/+^*, n=15). Data are mean ± SEM and box-and-whisker plots (medians with interquartile range, minimum, and maximum values are represented). \**P* < 0.05, \*\**P* < 0.01, \*\*\**P* < 0.001; (A and B) two-way ANOVA, (C) unpaired t-test, (D) two-way ANOVA with repeated measures, (E) two-way ANOVA with repeated measures in PPI test and Mann-Whitney U-test in startle response.

We performed the reciprocal dyadic social interaction test to evaluate social behavior, in which mice were allowed to freely move and reciprocally interact with each other (Harper et al., 2012; Hiramoto et al., 2011). *Auts2^del8/+^* mice displayed lower levels of active affiliative social interaction than WT mice in both session 1 and session 2 (Figure 6A). Of note, the restricted ablation of *Auts2* in mature excitatory neurons in the forebrain well recapitulated the impairment of social interaction, as depicted by tamoxifen-treated *CaMKIIa-CreER^T2^;Auts2^flox/flox^* mutants (Supplementary Figure 5A and D). Furthermore, in a three-chamber social interaction test, *Auts2^del8/+^* mutant mice displayed a decreased preference for a social subject (stranger mice 1 and 2) over non-social subject (empty chamber or familiar mouse) compared with WT mice in both sociability and social novelty phases (Figure 6B). These results suggest that *Auts2* mutant mice have social defects. We confirmed that sensory abilities such as olfaction and visual functioning as well as tactile response were not significantly different across the genotypes, as no phenotype was observed in the buried food finding test (Supplementary Figure 4A and 5C), whisker twitch reflex (100% response in WT, n=12, *Auts2^del8/+^*, n=10, *Auts2^flox/flox^*, n=10 and *CaMKIIa-CreER^T2^;Auts2^flox/flox^*, n=10) and visual placing response test (Supplementary Figure 4B and 5B), respectively. To further examine the sensory function of the vibrissae, we measured thigmotactic behaviors, defined as movement along the walls so that one side of the vibrissae could contact and scan the edge of the wall (Luhmann et al., 2005; Milani et al., 1989). *Auts2^del8/+^* mutant and WT mice behaved similarly in this test (Supplementary Figure 6C). These results suggest that the impaired social interaction probably does not involve the alterations in non-specific elements of social behavior such as sensory functioning.

Spontaneous locomotor activity test showed that the *Auts2^del8/+^* mice exhibited significantly decreased exploratory behavior during the first 15 min of the test (Supplementary Figure 4D).

In the open field test, the time that *Auts2^del8/+^* mice spent in the illuminated inner area was comparable to that of WT mice although general locomotor activity was slightly reduced in *Auts2^del8/+^* mice as indicated by total travel distance during the test (Supplementary Figure 4E). In the elevated plus maze test, however, *Auts2^del8/+^* mice displayed increased exploratory behavior of the open arms compared with WT mice, suggesting that *Auts2^del8/+^* mice have reduced fear of height (Figure 6C).

In a novel object recognition test, *Auts2^del8/+^* mice exhibited impaired recognition memory performance depicted by the significant decrease of time for exploratory index to the novel object (Figure 6D). Meanwhile, *Auts2^del8/+^* mice showed normal associative memory functions in the fear-conditioning test (Supplementary Figure 4F). Interestingly, *Auts2^del8/+^* exhibited a higher response to nociceptive stimuli as observed in the *Auts2^neo/+^* mutants in our previous study (Hori et al., 2015) (Supplementary Figure 4F). Furthermore, *Auts2^del8/+^* exhibited abnormal acoustic startle responses as well as sensorimotor gating deficits as indicated by decrease in the percentage of prepulse inhibition (Figure 6E).

### Altered vocal communication in *Auts2* mutant mice

Among types of social behaviors, mouse vocal communication has recently received attention as a possible model for studying the genetic and neural mechanisms for social communication (Holy and Guo, 2005). Mice use ultrasonic vocalizations (USVs) to exchange information in a variety of social contexts (Portfors and Perkel, 2014). When interacting with females, adult WT males actively emit courtship USVs with key tone frequencies between 50-80 kHz, as observed in the real-time spectrograms in Figure 5A. In contrast, the USVs produced by *Auts2^del8/+^* males were apparently dispersive during the test (Figure 7A). Indeed, the mean number and duration of USVs were markedly reduced in *Auts2^del8/+^* mice compared with WT controls (Figure 7B). Similarly, *CaMKIIa-CreER^T2^;Auts2^flox/flox^* males also displayed the altered vocalizations (Supplementary Figure 5E). The experiments of auditory playback previously showed that adult females prefer USVs with greater complexity from neonates as well as adult males (Chabout et al., 2015; Takahashi et al., 2016). We classified the acoustic structures of USVs into 12 different call patterns and grouped them into “simple” and “complicated” syllable types (Figure 7C). *Auts2^del8/+^* emitted significantly fewer numbers of the complicated syllable type, including “harmonics”, “complex” or “one jump + harmonics” whereas the simple syllable types with shorter duration such as “downward” or “short” were significantly increased (Figure 7D). These findings suggest that loss of *Auts2* alters mouse vocal communication, which may underlie the pathology for communication disorders in ASD patients with *AUTS2* mutations.

**Figure 7.**
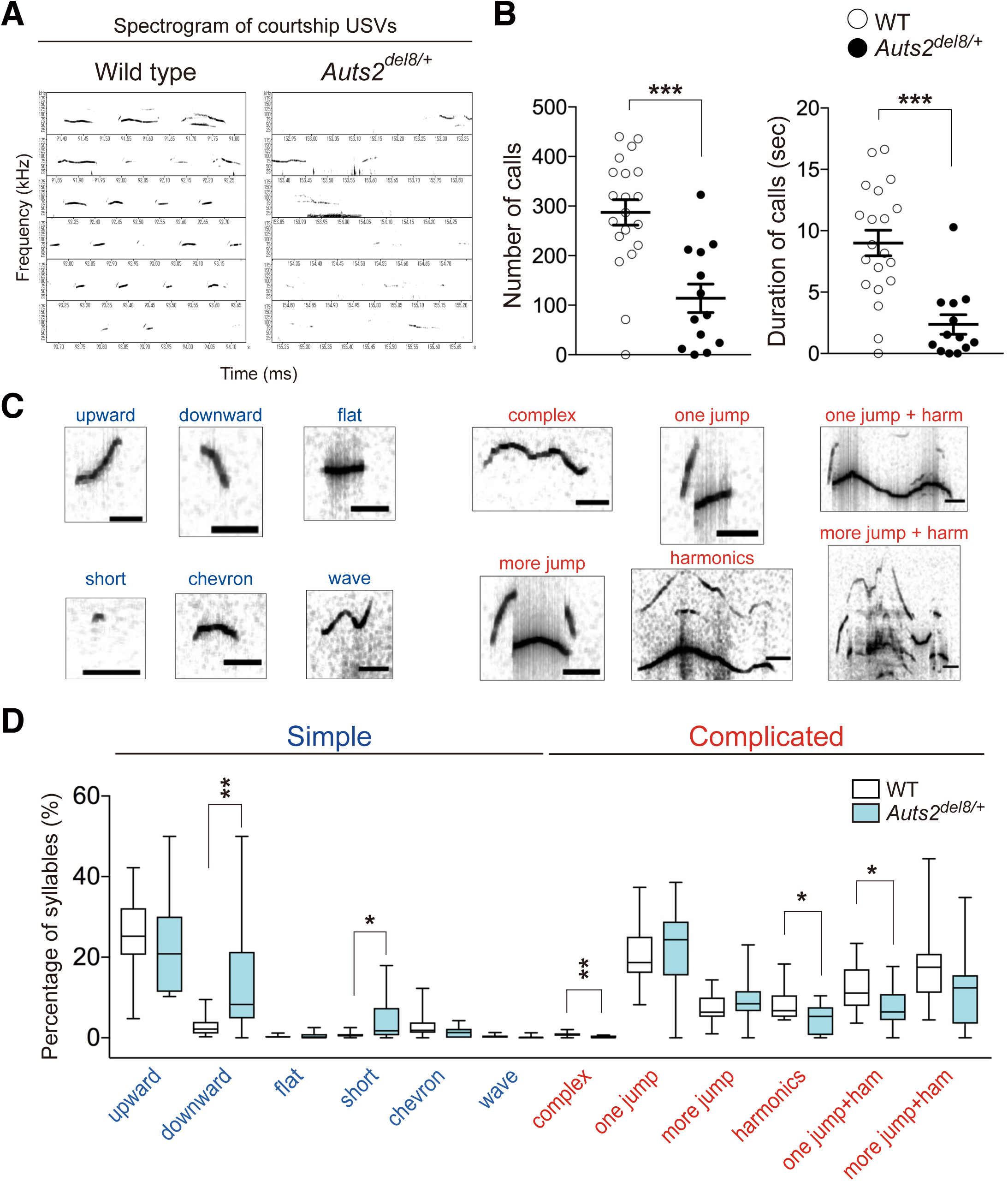
Deficits in vocal communication in adult *Auts2^del8^* mutant mice. (A) Representative spectrograms of USV during the courtship behaviors. (B) The number (left) and duration (right) of USVs during 1 min. (C) Typical spectrograms of 12 different call patterns. Six simple call types (blue) and six complicated call types (red) are indicated. (D) The frequency of each syllable pattern is shown as the percentage of total calls. Data are mean ± SEM and box-and-whisker plots (medians with interquartile range, minimum, and maximum values are represented). (WT, n=20, *Auts2^del8/+^*, n=13). \**P* < 0.05, \*\**P* < 0.01, \*\*\**P* < 0.001; (B) unpaired t-test, (D) Mann-Whitney U test.

## [Discussion]

In this study, we found that AUTS2 restricts the number of excitatory synapse during development as well as in adults. In *Auts2* mutant brains, excessive numbers of dendritic spines are formed on the pyramidal forebrain neurons at several brain areas including prefrontal and temporal cortex as well as hippocampus, which are implicated as the critical regions for cognitive brain functions. Interestingly, in *Auts2* mutant cerebral cortex, aberrant spine formation specifically appeared in the upper-layer but not deep-layer neurons although AUTS2 is widely expressed in both cortical layers (Figure 2B and Supplementary Figure 2D) (Bedogni et al., 2010). One plausible hypothesis is that AUTS2 may have distinct roles for neural development in different cerebral cortical areas, which may depend on differences of AUTS2 isoforms expressed between neurons or on co-factors that differentially interact with each AUTS2 isoform. Monderer-Rothkoff et al have recently demonstrated that the long and short AUTS2 isoforms, each interacting with different co-factors, act opposingly on gene transcription in a cellular-context dependent manner (Monderer-Rothkoff et al., 2019).

Electrophysiological experiments revealed that excitatory but not inhibitory synaptic inputs were elevated in the *Auts2* mutant hippocampal slices where strong c-Fos signals were observed, implying that a balance between excitation and inhibition (E/I) was disturbed in that region. E/I balance in neural circuits is tightly controlled and established by contributions from a large number of factors in the normal brain. Accumulating evidence implicates a disturbed E/I balance within cortical neural circuitry in various neuropsychiatric disorders including ASD, anxiety and ADHD (Chao et al., 2010; Edden et al., 2012; Gogolla et al., 2009; Han et al., 2012; Rubenstein and Merzenich, 2003). Although a recent report suggests that E/I imbalance is not causative for the neuropathology of the disorders but reflects a homeostatic response in some mouse models (Antoine et al., 2019), the hyperexcitability caused by an increased E/I ratio in the cerebral cortex is thought to be one potential common mechanism underlying the neurobehavioral defects of some forms of ASD via a distinct molecular pathway (Lee et al., 2017).

During the spinogenesis, a rapid increase of dendritic spine density occurs in the forebrain neurons, in which the gain of spines exceeds loss of spines, eventually causing excessive excitatory synapses for the formation of neural circuits (Chen et al., 2014; Forrest et al., 2018; Isshiki et al., 2014; Penzes et al., 2011). Thereafter, the growth of excitatory synapses is gradually downregulated and unnecessary spines are selectively pruned, after which spines are maintained during adulthood. Time-lapse imaging experiments using *Auts2*-knocked-down hippocampal neurons revealed that *de novo* formation of dendritic spines is promoted whereas the elimination rate is decreased, resulting in the exaggerated formation of excitatory synapses. These observations suggest an important role for AUTS2 in controlling the number of spines or excitatory synapses in forebrain neurons by modulating their turnover. We found that this excess in synapses was also observed in tamoxifen-treated *CaMKIIa-CreERT2;Auts2^flox/flox^* in which *Auts2* was ablated after establishment of the brain structure. This suggests that AUTS2 is involved in regulating synaptic homeostasis at late developmental and/or adult stages.

Emerging evidence indicates that aberrant regulation of spine number and/or an increased excitatory synaptic inputs likely caused by incomplete pruning or exaggerated formation of spines is associated with numerous pathological conditions such as ASD, schizophrenia and neurodegenerative disorders (Chen et al., 2014; Forrest et al., 2018; Lee et al., 2017; Penzes et al., 2011). Transcriptional control by epigenetic regulation including histone post-translational modification and chromatin remodeling is critical in synapse development and neurological disorders. A recent study by Korb et al. revealed that Fragile X mental retardation protein *Fmr1* mutant mice exhibit widespread histone mis-modifications (Korb et al., 2017). These are associated with open chromatin caused by upregulation of epigenetic factor Brd4, resulting in alteration of the transcription levels of many critical synapse-related genes. In this study, we showed that nuclear-localizing AUTS2 functions to restrict spine number. Because AUTS2 is involved in transcriptional regulation via chromatin modification as a component of PRC1 (Gao et al., 2014), and because expression of many synapse-related genes were altered in the *Auts2* mutants (Figure 5), we believe that nuclear AUTS2 restricts the excitatory synapse number via controlling the expression of relevant genes, thus maintaining the excitation/inhibition balance of the brain.

In previous and current studies, we characterized behavioral phenotypes for two lines of mutant mice with different mutations disrupting the *Auts2* locus (Hori et al., 2015). We summarized the results from a behavioral test battery for *Auts2^neo/+^* (Hori et al., 2015) and *Auts2^del8/+^* mutant mice (this study) in Supplementary Figure 6G. In this study, we found that the *Auts2^del8/+^* heterozygous global KO as well as *CaMKIIa-CreER^T2^;Auts2^flox/flox^* conditional KO mice exhibited autistic-like behaviors including social deficits and altered vocal communications as well as multiple other behavioral impairments.

In addition, *Auts2^del8/+^* mice also showed altered anxiety as well as higher responses against nociceptive and auditory stimuli, both of which are often observed in ASD patients (American Psychiatric Association, 2013). Interestingly, *Auts2^del8/+^* mutant mice share several behavioral phenotypes with *Auts2^neo/+^* mutants but also display a distinct combination of phenotypes (Supplementary figure 4G). Although the underlying mechanisms how different mutations lead to the distinct behavioral phenotypes in mice remains unclear, it is possible that compensatory expression of an AUTS2 C-terminal short isoform in *Auts2^del8/+^* mutant brains negatively affects social behaviors in the social interaction tests (Figure 6A and B), whereas it alleviates the cognitive dysfunctions displayed in *Auts2^neo/+^* mutant mice (Hori et al., 2015) such as the associative memory formation in fear-conditioning tests (Supplementary Figure 4F).

In humans, it has been reported that multiple types of heterozygous genomic structural variants in the *AUTS2* locus including *de novo* balanced translocation, inversion or intragenic deletions are associated with a wide range of psychiatric illnesses such as ASDs, ID, ADHD, schizophrenia and dyslexia as well as other neuropsychiatric diseases (Oksenberg and Ahituv, 2013). In addition to the exonic deletions of the *AUTS2* locus, some of the genomic structural variants are within intronic regions, implying that improper and disorganized expression of *AUTS2* could be involved in the onset of the disorders. However, it remains largely unclear how different mutations of the same gene contribute to different diseases. Currently, eight computationally annotated *AUTS2* isoforms in humans are incorporated in public databases (for example: the UCSC Genome Bioinformatics (https://genome.ucsc.edu)). However, the study by Kondrychyn et al. revealed that *auts2a*, the zebrafish ortholog of *Auts2*, possesses 13 putative unique transcriptional start sites (TTS) and surprisingly, more than 20 alternative transcripts are potentially produced from this gene locus by the aforementioned TSSs and/or by alternative splicing (Kondrychyn et al., 2017). These findings suggest that mammals including mouse and human could have similar or higher transcriptional complexity for *Auts2*/*AUTS2* than previously thought. Furthermore, Oksenberg et al. have identified several enhancer regions for the expression of *auts2a/Auts2* in zebrafish and mouse brain within the intronic regions of this gene locus (Oksenberg et al., 2013). Therefore, structural variants such as genomic deletions within a certain region of *Auts2/AUTS2* locus could not only alter the expression of full-length AUTS2 directly, but also affect the transcriptional regulation of other AUTS2 isoforms. Different mutations of the *AUTS2* gene may differentially alter the temporal and spatial expression profiles of AUTS2 isoforms in various brain regions, which may distinctively affect neurobiological functions, ultimately resulting in the occurrence of multiple types of psychiatric disorders in individuals with *AUTS2* mutations. Our previous and this study, thus, highlighted that two types of *Auts2* mutants with different AUTS2 protein expression profiles exhibited overlapping but distinct behavioral abnormalities. This may support the notion that different types of mutations in *AUTS2* account for distinct types of neuropsychiatric illnesses. Future comprehensive studies elucidating the regulatory mechanisms for transcription/splicing of *Auts2/AUTS2* as well as neurobiological functions of the distinctive AUTS2 isoforms will help us to understand the pathogenic mechanisms underlying the occurrence of a variety of psychiatric disorders in individuals with *AUTS2* mutations, and could contribute to therapeutic development for *AUTS2*-related neurological disorders.

In conclusion, the findings presented here suggest that synaptic regulation by AUTS2 is required for proper social behaviors. Furthermore, our results from the behavioral analyses for *Auts2^del/8/+^* KO mice provided insight into the involvement of AUTS2 in other higher brain functions such as recognition and emotion. In addition to the AUTS2 function on synapse regulation, AUTS2 is also involved in neuronal migration and neurite formation (Hori et al., 2014). Therefore, the other abnormal behaviors observed in *Auts2^del/8/+^* or *Auts2^neo/+^* KO mice may partly be caused by the impairments in these developmental processes. Comparative analyses of the different forms of *Auts2* mouse mutants will help us to better understand the pathological mechanisms of the psychiatric disorders caused by *AUTS2* mutations. *Auts2* conditional KO mice with *CaMKIIa-CreER^T2^* or other more restricted-expression forms of *Cre* will be useful for dissecting the distinct neural circuitries involved in these abnormal behaviors.

## [Methods]

### Experimental animals

*Rosa26R^YFP^* mouse line (stock no. 006148) was obtained from The Jackson Laboratory. Generation of *Auts2^del8^* and *Auts2-floxed* mice with a pure C57BL/6N genetic background and genotyping for these mice have been previously described (Supplementary Table 1) (Hori et al., 2014). *Emx1^Cre^* (stock #RBRC00808, C57BL/6J background) and *CaMKIIa-CreER^T2^* (B6.FVB-Tg(Camk2a-cre/ERT2)2Gsc/Ieg, stock #EM02125) mice were purchased from RIKEN BioResource Center (RIKEN, Tsukuba, Japan) and European Mouse Mutant Archive (EMMA) (HelmholtzZentrum München, Neuherberg, Germany), respectively (Erdmann et al., 2007; Iwasato et al., 2000). For the experiments with *CaMKIIa-CreER^T2^* mice, tamoxifen (Sigma-Aldrich, St. Louis, MO, USA) was administered at 50 mg/kg during postnatal 21-25 days for anatomical analysis or P30-34 for behavioral analyses by intraperitoneal injection twice daily for 5 consecutive days and the analyses were performed at postnatal day 50 and 10-12 weeks, respectively. Mice were maintained in ventilated racks under a 12-h light/dark cycle with food and water ad libitum in temperature controlled, pathogen-free facilities. Mice of each genotype were randomly allocated to different experiments.

### Behavioral analysis

For behavioral test battery using *Auts2^del8^* mice, two independent cohorts of *Auts2^del8/+^* heterozygotes and their wild type littermate male mice were tested, to confirm findings. All behavioral tests using *Auts2^del8^* mice were obtained by crossing *Auts2^del8/+^* heterozygous male mice with wild type C57BL6/N female mice (Charles River Laboratories, Kanagawa, Japan) to avoid the possibility that altered behaviors in the mutant dams could influence the postnatal development of their offspring. After weaning, male mice were cohoused in same-genotype groups of 2-4 littermates per cage before and during the behavioral tests.

Behavioral tests were performed using the same set of mice in the following sequence: locomotor activity, open field test, novel object recognition test, elevated plus maze, 3-chamber social interaction test, prepulse inhibition test, and fear conditioning test. For recording of USVs, buried food finding test, visual placing response test, thigmotaxis and reciprocal social interaction test, separate cohorts of mice were used.

For behavioral analyses using *CaMKIIa-CreER^T2^;Auts2^flox^* conditional KO mice, *CaMKIIa-CreER^T2^;Auts2^flox/flox^* homozygous mutant mice and their control littermate *Auts2^flox/flox^* male mice were used.

### Plasmid construction

The plasmid construction of pCAG-Myc-AUTS2-full length, FL-AUTS2^R^ (the shRNA-resistant AUTS2-full length), NES-FL-AUTS2^R^ and S-AUTS2-var.2 were previously described (Hori et al., 2014). cDNA fragments for S-AUTS2-var.1 encoding 1,372-3,786 bp were amplified by PCR using full-length *Auts2* cDNA as a template and subcloned into pCAGGS vector. To construct the 3xNLS-AUTS2 expression plasmid, two oligonucleotides coding the three tandem nuclear localization signal (NLS) sequence of the SV40 Large T-antigen (PKKKRKV) were annealed and inserted between Myc-tag and 5’-terminus of AUTS2 ORF in pCAG-Myc-AUTS2^R^ with EcoRI site: Fwd, (5’-AATTGGTGCACGTGGATCCAAAAAAGAAGAGAAAGGTAGATCCAAAAAAGAAGAGAAAGGTAGATCCAAAAAAGAAGAGAAAGGTACACGTGTCCG-3’): Rev, (5’-AATTCGGACACGTGTACCTTTCTCTTCTTTTTTGGATCTACCTTTCTCTTCTTTTTTGGATCTACCTTTCTCTTCTTTTTTGGATCCACGTGCACC-3’). The expression plasmids for EGFP, Cre recombinase and shRNAs were described previously (Hori et al., 2014).

### Primary culture of hippocampal neurons

Primary hippocampal cultures were prepared as previously described with minor modifications (Hori et al., 2014; Hori et al., 2005). Hippocampi at E17.5 were dissected from C57BL6/N wild type or homozygotic *Auts^2flox/flox^* mouse embryos and dissociated using Neuron Dissociation Solution S (Fujifilm Wako Pure Chemical Corporation, Osaka, Japan). The dispersed neurons were electroporated with the expression plasmids or shRNA vectors using the NEPA21 electroporator (Nepa Gene, Chiba, Japan) according to the manufacturer’s instructions. The electroporated neurons were mixed with the transfection-free control neurons at a ratio of approximately 1:20 and plated on coverslips coated with 0.1 mg/ml poly-D-lysine (Sigma-Aldrich, St. Louis, MO, USA) at a density of 8,000 – 12,000 cells/cm^2^ and maintained in astroglial-conditioned Neurobasal medium containing 2% B27 supplement (Thermo Fisher Scientific, Waltham, MA, USA) and 1 mM L-glutamine.

### Immunostaining

For immunocytochemistry, cells were fixed with 4% PFA/4%sucrose for 40 min on ice. Immunostaining was performed using the following primary antibodies: mouse anti-PSD-95 (6G6-1C9, ThermoFisher Scientific, Waltham, MA, USA), mouse anti-Gephyrin (3B11, Synaptic Systems, Goettingen, Germany), rat anti-GFP (RQ1, gift from A. Imura, BRI, Kobe). For immunohistochemistry, adult mouse brains were dissected after mice were deeply anesthetized and transcardially perfused with 4% PFA. The brains were post-fixed with 4% PFA/5%sucrose for 6 hrs or overnight at 4 °C, rinsed with PBS, cryoprotected with 30% sucrose, embedded in O.C.T compound (Sakura Fine-Tek, Tokyo, Japan), and cryosectioned at 14∼30 μm. For c-Fos staining, tissue sections were immunostained with rabbit anti-c-Fos antibody (sc-52, Santa Cruz Biotechnology, Inc., Dallas, TX, USA). Acquisition of fluorescent images, counts and measurement of dendritic protrusions were carried out using a Zeiss LSM 780 confocal microscope system and ZEN software (Carl Zeiss, Oberkochen, Germany). For DAB staining with rat anti-GFP antibody (RQ1), the sections were processed using the VECTASTAIN ABC system (Vector Laboratories, Burlingame, CA, USA) with diaminobenzidine (Sigma-Aldrich, St. Louis, MO, USA) and images were taken on Keyence All-in-One fluorescence microscope (BZ-X700, Osaka, Japan).

### Golgi-staining and analysis of dendritic spines

Whole brains collected from mice were subjected to Golgi impregnation solution (FD Rapid GolgiStain kit, FD NeuroTechnologies, Columbia MD, USA). Coronal sections with 80-100 μm thick were obtained with cryostat and mounted on gelatin-coated slides. After tissues were processed for Golgi-Cox staining according to manufacturer’s instructions, the brain sections were dehydrated with a graded series of ethanol, immersed in xylene, and embedded in Entellan (Merck, Darmstadt, Germany). Neurons were traced using a Leica microscope (DM5000B, Leica Microsystems, Danaher, Germany) with a 100x oil-immersion objective and were 3D-reconstructed by Neurolucida Software (MBF Bioscience, Williston, VT, USA). Spines along the dendritic segments (>30 μm length) within 100 μm from soma were examined, and spine densities were calculated as mean number of spines per 10 μm dendrites. On the basis of spine morphology, dendritic protrusions were classified into the following four categories (Harris and Kater, 1994) : thin (≥ 0.5 μm protrusions with small bulbous head less than twice as large as spine neck), mushroom (≥0.5 μm protrusions with bulbous head more than twice as large as spine neck), stubby (≥0.5 μm protrusions with bulbous head but without a neck), filopodia (≥5 μm long and thin protrusions without bulbous heads). Dendritic protrusions with total lengths exceeding 10 μm were considered as branched dendrites and excluded from the analysis.

### Electrophysiology

Whole-cell voltage-clamp recordings of mEPSCs and mIPSCs using brain slices were conducted as previously described (Takahashi et al., 2012). Coronal hippocampal slices with 400 μm thickness from adult mice at P33-44 were prepared in ice-cold dissection buffer (300 mM sucrose, 3.4 mM KCl, 0.6 mM NaH_2_PO_4_, 10 mM D-Glucose, 10 mM HEPES, 3.0 mM MgCl_2_, 0.3 mM CaCl_2_ at pH 7.4) using a VT1200S vibratome (Leica Biosystems, Danaher, Germany). Hippocampal slices were incubated in artificial cerebrospinal fluid (ACSF; 119 mM NaCl, 2.5 mM KCl, 1.0 mM NaH_2_PO_4_, 26.2 mM NaHCO_3_, 11 mM D-Glucose, 4.0 mM MgSO_4_, 4.0 mM CaCl_2_, gassed with 95% O_2_ and 5% CO_2_), left to recover for more than 1 hour at room temperature, and then transferred to a recording chamber mounted on an upright microscope (BX61WI, Olympus, Tokyo, Japan). For voltage-clamp recordings of hippocampal slices, borosilicate glass pipettes (4-6 MΩ) were filled with the internal solutions (135 mM CsMeSO_4_, 10 mM HEPES, 0.2 mM EGTA, 8 mM NaCl, 4 mM Mg-ATP, 0.3 mM Na_3_GTP at pH7.2, osmolality adjusted to 280-300 mOsm). All data of whole-cell voltage-clamp recordings were acquired with Multiclamp 700B (Molecular Devices, San Jose, CA, USA) equipped with an A/D converter (BNC-2090, National Instruments, Austin, TX, USA) and Igor Pro software version 4.01 (Wavemetrics, Portland, OR, USA) at 4 kHz. Series resistances were monitored, and the data were discarded when the series resistance changed by > 30 MΩ during recordings. mEPSCs were recorded at −70 mV in the presence of 1 μM tetrodotoxin and 100 μM picrotoxin, and mIPSCs were recorded at 0 mV in the presence of 1 μM tetrodotoxin, 10 μM CNQX and 50 μM D-APV. mEPSCs and mIPSCs events above a threshold value (10 pA) were analyzed with Minianalysis software version 6.0.3 (Synaptosoft, Fort Lee, NJ, USA).

### Sample size and statistical analysis

Sample size was determined based on studies using established methods and on our previous experiments (Hori et al., 2015; Hori et al., 2014; Hori et al., 2005; Takahashi et al., 2012). Data analyses were performed blinded to the genotype. The number of samples and animals is indicated in the figure legends. All statistical analyses except transcriptome data processing and analysis were performed using GraphPad Prism 7 (GraphPad Software, La Jolla CA, USA). The normal distribution of data was confirmed by the Shapiro-Wilks test and if significant, a nonparametric Mann Whitney U test was used for comparison. Equal variance was tested by the F-test and when there was a significant difference, we used a two-tailed unpaired t-test with Welch’s correction. When the data were within the assumptions of normal distribution and equal variance, a two-tailed unpaired t-test was used for comparison of the means between two groups. For comparison of more than 2 groups, a one-way analysis of variance (ANOVA) followed by the Dunnett’s multiple comparison test was used.

In the behavioral analysis, two-way ANOVA followed by the Bonferroni test was used for multiple-group comparisons (reciprocal social interaction test and three-chamber social interaction test). Two-way ANOVA with repeated measurements followed by the Bonferroni test was used for multiple-group comparisons (locomotor activity, novel object recognition test, thigmotaxis and prepulse inhibition test).

## Author contributions

KH designed this study. KH and MH wrote the manuscript and coordinated the project. KH, MY, SE, KS, AS and MS performed and supervised imaging experiments and statistical analysis; WS, TN, and AS carried out, and KY supervised behavioral experiments and data analysis; KY, KS and HK performed and supervised electrophysiological experiments; RS and KK performed and supervised recording and analysis of ultrasonic vocalizations; MA and KS generated and supervised the designs of *Auts2* mutant mice. YG, ST and SM performed RNA-seq and data analysis.

## Acknowledgement

This work was supported by Grants-in-Aid for Scientific Research, KAKENHI (Grant 16H06528 and 18H02538 to M.H. and 16K07021 to K.H.), and Innovative Areas (16H06524 and 16H06531 to Y.G.) from MEXT; the SRPBS from AMED (19dm0107085h0004), Naito Foundation, Takeda Foundation, Uehara Foundation, Suzuken Memorial Foundation, Princess Takamatsu Cancer Research Fund, an Intramural Research Grant (Grants 30-9 and 1-4 to M.H.). We are grateful to Dr. Ruth Yu (St Jude Children’s Research Hospital) for comments on the manuscript.

## Competing interests statement

The authors have declared that no conflict of interest exists.

## Notes

https://www.ncbi.nlm.nih.gov/geo/query/acc.cgi?acc=GSE134712

