## Supplemental figures and methods for "AUTS2 regulation of synapses for proper synaptic inputs and social communication"

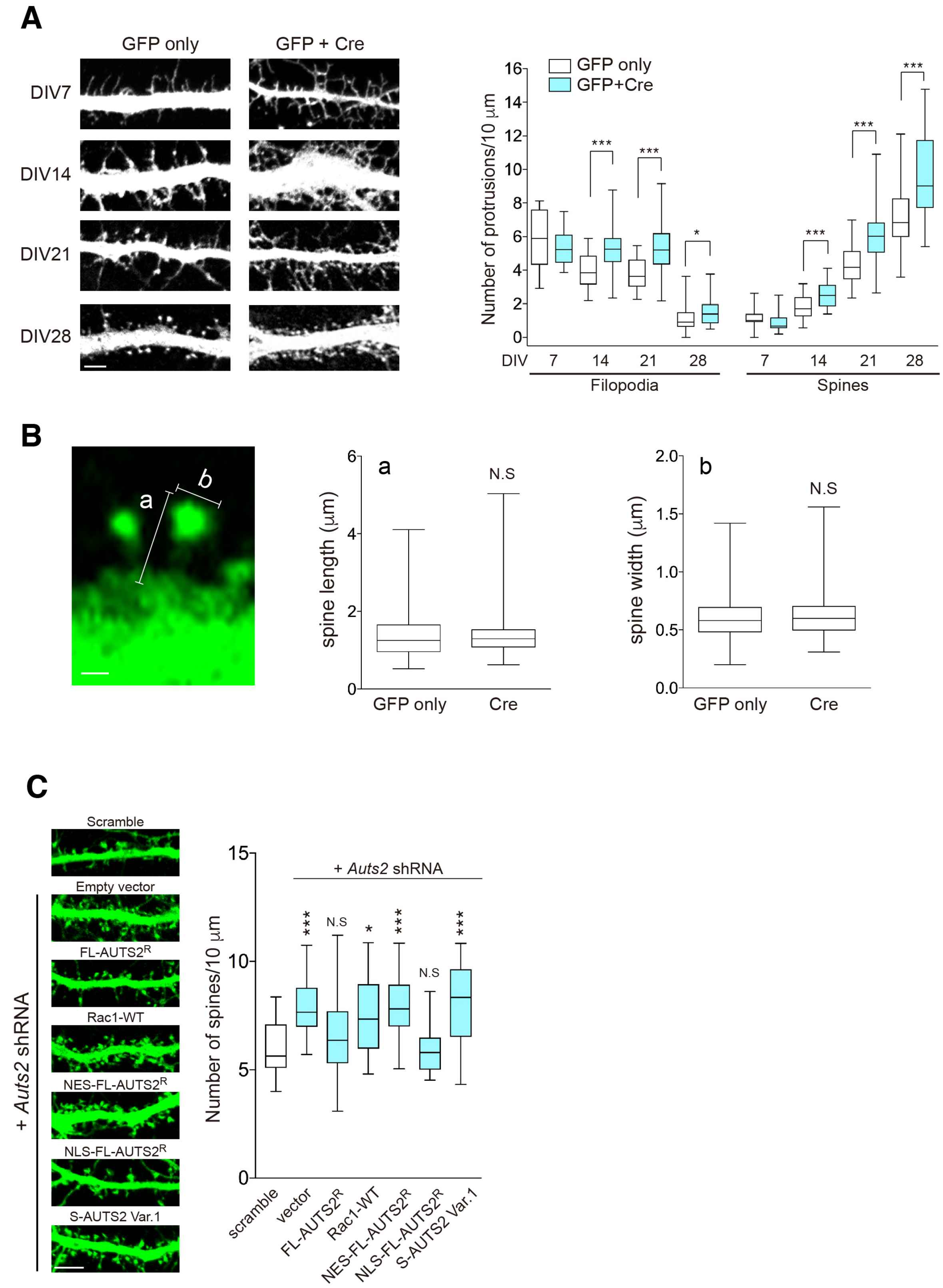
A
GFP only
GFP + Cre
DIV7
DIV14
DIV21
DIV28
GFP only
GFP+Cre
Number of protrusions/10 μm
16
14
12
10
8
6
4
2
0
***
***
*
***
***
***
DIV
7
14
21
28
7
14
21
28
Filopodia
Spines
B
a
b
6
4
2
0
spine length (μm)
N.S
GFP only
Cre
2.0
1.5
1.0
0.5
0.0
spine width (μm)
N.S
GFP only
Cre
C
Scramble
Empty vector
FL-AUTS2R
Rac1-WT
NES-FL-AUTS2R
NLS-FL-AUTS2R
S-AUTS2 Var.1
+ Auts2 shRNA
+ Auts2 shRNA
15
10
5
0
Number of spines/10 μm
***
N.S
*
***
N.S
***
scramble
vector
FL-AUTS2R
Rac1-WT
NES-FL-AUTS2R
NLS-FL-AUTS2R
S-AUTS2 Var.1

**Supplementary Figure 1.** ***Auts2*** **mutant primary neurons exhibit excessive spine formation. Related to Figure 1**

**(A)** Primary hippocampal neurons derived from *Auts2^flox/flox^* homozygotes were electroporated with the control or Cre expression vectors. To visualize the neurons, GFP expression vector was co-electroporated. The density of dendritic spines and filopodia on the dendrites of the control and *Auts2* KO (Cre) neurons were measured at different culture stages (DIV7-28) (n=21-40 dendrites of 11-20 neurons). **(B)** The length of dendritic spines (a) and width of the spine head (b) in the control and *Auts2* KO neurons (Cre) at DIV28 were measured (control; n=155 spines, Cre; n=158 spines). **(C)** WT primary hippocampal neurons were co-electroporated with *Auts2*-shRNA and the indicated expression vectors. To visualize the neurons, GFP vector was co-electroporated. Graph shows the density of dendritic spines (n = 20 dendrites). Expression of the shRNA-resistant FL-AUTS2 (FL-AUTS2$^{R}$) or nuclear-localized form AUTS2 (NLS-AUTS2$^{R}$) in *Auts2*-knockdown neurons rescues the aberrant spine formation. Data are mean ± SEM and box-and-whisker plots (medians with interquartile range, minimum, and maximum values are represented). $*P < 0.05$, $***P < 0.001$, N.S, not significant. (A) unpaired t-test, (B) Mann-Whitney U test, (C) one-way ANOVA with Dunnett's post hoc test. Quantifications represent data from three independent experiments. Scale bar, 10 μm in (A and C) and 2 μm in (B).

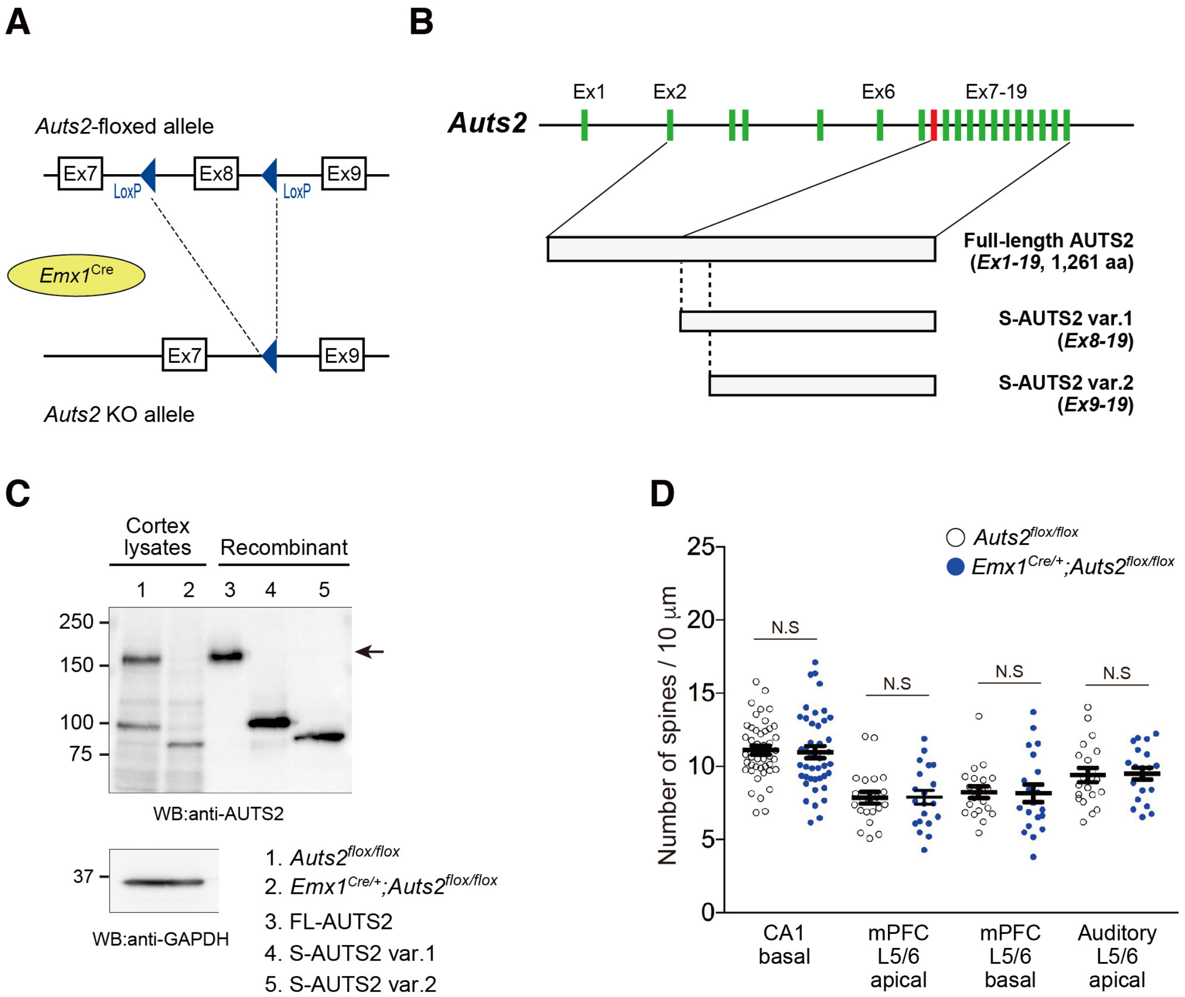
A
Auts2-floxed allele
Ex7
Ex8
Ex9
LoxP
Emx1Cre
Auts2 KO allele
B
Auts2
Ex1
Ex2
Ex6
Ex7-19
Full-length AUTS2
(Ex1-19, 1,261 aa)
S-AUTS2 var.1
(Ex8-19)
S-AUTS2 var.2
(Ex9-19)
C
Cortex lysates
Recombinant
1 2 3 4 5
250
150
100
75
WB:anti-AUTS2
37
WB:anti-GAPDH
1. Auts2flox/flox
2. Emx1Cre/+;Auts2flox/flox
3. FL-AUTS2
4. S-AUTS2 var.1
5. S-AUTS2 var.2
D
Auts2flox/flox
Emx1Cre/+;Auts2flox/flox
Number of spines / 10 μm
25
20
15
10
5
0
N.S
CA1 basal
mPFC L5/6 apical
mPFC L5/6 basal
Auditory L5/6 apical

**Supplementary Figure 2. Analysis of spine formation in forebrain-specific *Auts2* conditional KO mice. Related to Figure 2**

**(A)** Schematics of the targeting strategy. The deletion of exon 8 at *Auts2* locus in pyramidal neurons of the forebrain was generated by crossing the *Auts2-floxed* mice with $Emx1^{Cre}$ mice. **(B)** Schematic of *Auts2* genomic region and the protein structure of AUTS2 isoforms. **(C)** Western blotting of lysates from P0 cerebral cortex of $Auts2^{flox/flox}$ (Control) and $Emx1^{Cre/+}$;$Auts2^{flox/flox}$ homozygotes using anti-AUTS2 antibody. Immunoblot of lysates from HEK293T cells expressing the recombinant full-length AUTS2 and the C-terminal AUTS2 short variants (S-AUTS2 var.1 and var.2) are also shown. Full-length AUTS2 (arrow) as well as the S-AUTS2 var.1 were completely eliminated in *Auts2* homozygotic mutant cerebral cortices whereas the S-AUTS2 var.2 was alternatively increased. **(C)** Summary graph of the spine density on the basal dendrites of CA1 pyramidal neurons, apical and basal dendrites of the deep-layer (L5/6) neurons at mPFC and auditory cortex in the $Auts2^{flox/flox}$ and $Emx1^{Cre/+}$;$Auts2^{flox/flox}$ homozygotic mutant mouse brains. (n=20 dendrites from N=3 animals) Data are presented as mean ± SEM. (D) unpaired t-test.

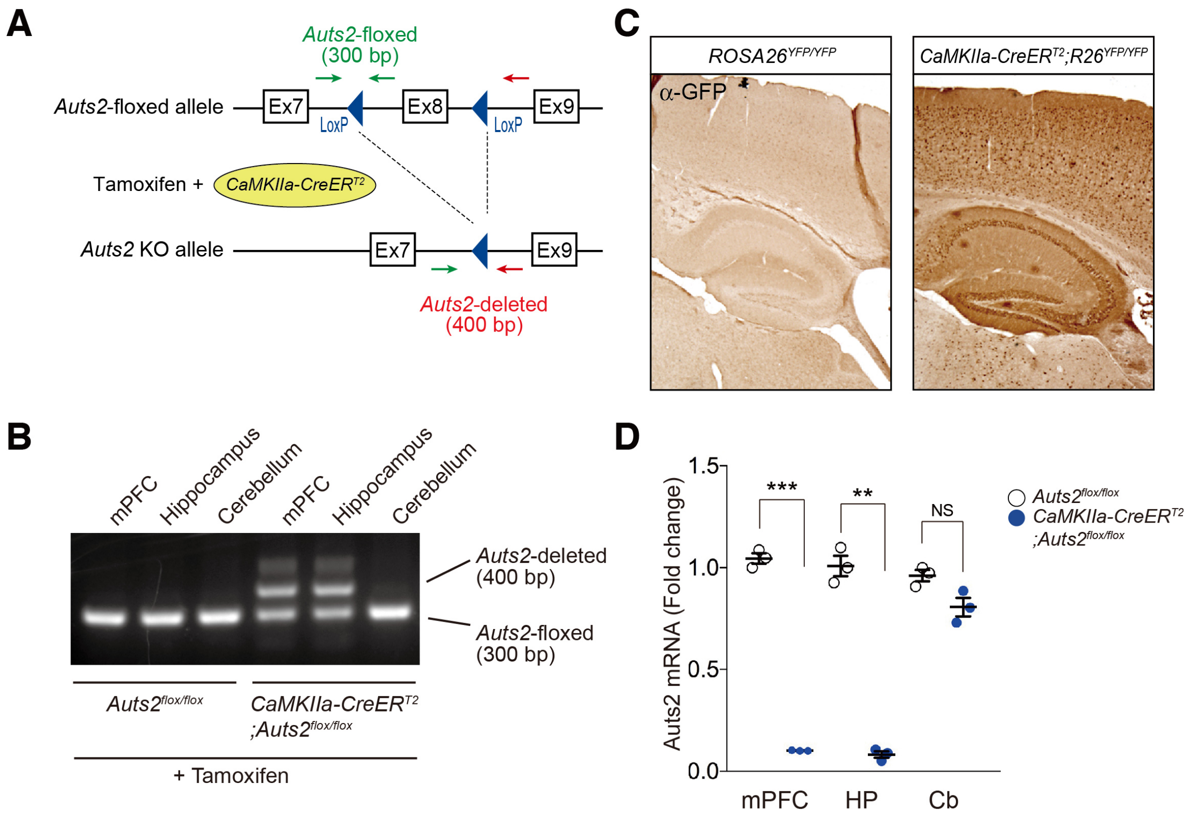

A
Auts2-floxed
(300 bp)
Auts2-floxed allele
Ex7
Ex8
Ex9
LoxP
LoxP
Tamoxifen +
CaMKIIa-CreERT2
Auts2 KO allele
Ex7
Ex9
Auts2-deleted
(400 bp)
B
mPFC
Hippocampus
Cerebellum
mPFC
Hippocampus
Cerebellum
Auts2-deleted
(400 bp)
Auts2-floxed
(300 bp)
Auts2flox/flox
CaMKIIa-CreERT2
;Auts2flox/flox
+ Tamoxifen
C
ROSA26YFP/YFP
CaMKIIa-CreERT2;R26YFP/YFP
α-GFP
D
Auts2 mRNA (Fold change)
1.5
1.0
0.5
0.0
***
**
NS
mPFC
HP
Cb
Auts2flox/flox
CaMKIIa-CreERT2
;Auts2flox/flox

**Supplementary Figure 3. Conditional deletion of *Auts2* in postnatal forebrain leads to excessive spine formation. Related to Figure 4**

**(A)** Schematics of the targeting strategy. The inducible conditional deletion of exon 8 at *Auts2* locus in pyramidal neurons of the forebrain was generated by breeding the *Auts2*$^{flox/flox}$ mice to *CaMKIIa-CreER*$^{T2}$ mice. **(B)** The forebrain-specific deletion of *Auts2* locus in the homozygous mutants (*CaMKIIa-CreER*$^{T2}$*;Auts2*$^{flox/flox}$) and control littermates (*Auts2*$^{flox/flox}$) after tamoxifen administration was confirmed by genomic PCR using the primer pairs indicated as green and red arrows in (A). **(C)** Examination of CreER$^{T2}$ recombinase activity in the postnatal brain. Tamoxifen was administered to *CaMKIIa-CreER*$^{T2}$*;ROSA26R*$^{YFP/YFP}$ reporter mice and the *ROSA26R*$^{YFP/YFP}$ control littermate during P21-25. Brain sections isolated 10 days after tamoxifen treatment were DAB-stained with anti-GFP antibody. The expression of EYFP was observed in the cortex and hippocampus in the presence of the *CreER*$^{T2}$ transgene (right panel). **(D)** Examination of *Auts2* transcript levels in mPFC, hippocampus (HP) and cerebellum (Cb) of adult *CaMKIIa-CreER*$^{T2}$*;Auts2*$^{flox/flox}$ homozygotic mutant mice and the control mice (*Auts2*$^{flox/flox}$) with tamoxifen application. qPCR was performed using primers specific for the deleted exon (n=3 brains). Data are mean ± SEM and box-and-whisker plots (medians with interquartile range, minimum, and maximum values are represented). $^{**}P < 0.01$, $^{***}P < 0.001$, unpaired t-test. Scale bar, 10 µm.

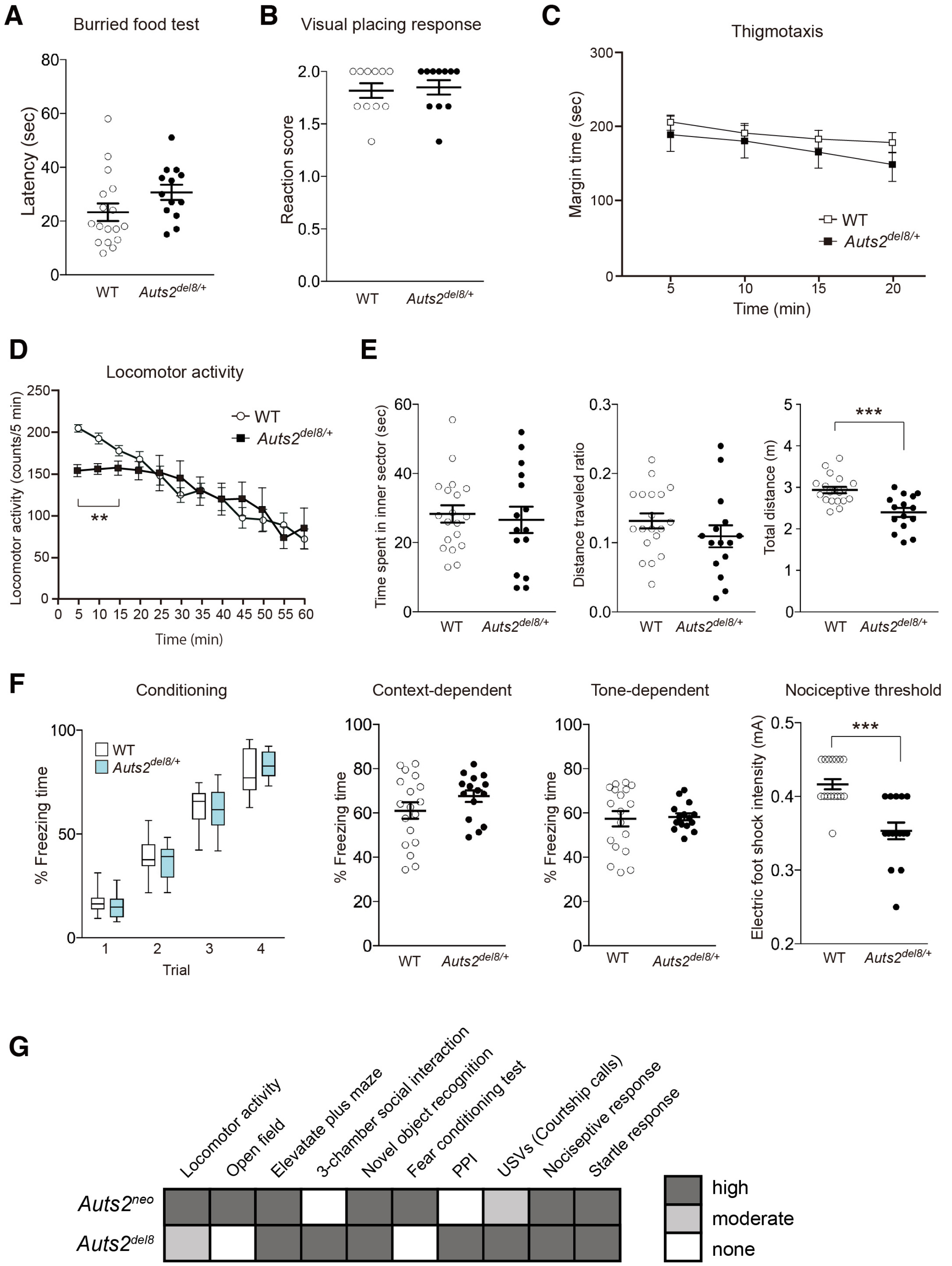

A
Burried food test
Latency (sec)
WT
Auts2del8/+
B
Visual placing response
Reaction score
C
Thigmotaxis
Margin time (sec)
Time (min)
D
Locomotor activity
Locomotor activity (counts/5 min)
Time (min)
E
Time spent in inner sector (sec)
Distance traveled ratio
Total distance (m)
F
Conditioning
% Freezing time
Trial
Context-dependent
Tone-dependent
Nociceptive threshold
Electric foot shock intensity (mA)
G
Locomotor activity
Open field
Elevatate plus maze
3-chamber social interaction
Novel object recognition
Fear conditioning test
PPI
USVs (Courtship calls)
Nociceptive response
Startle response
Auts2neo
Auts2del8
high
moderate
none

**Supplementary Figure 4. Behavioral analyses of *Auts2*$^{del8/+}$ mutant mice. Related to Figure 6**

**(A)** The buried food finding test. Time spent to find buried food pellet was measured. (WT, n=17, *Auts2*$^{del8/+}$ n=13). **(B)** Visual placing response test. Reaction score was rated as follow: 0, no observable placing behavior; 1, a weak or delayed placing response; 2, a clear placing reaction. (WT, n=11, *Auts2*$^{del8/+}$ n=11). **(C)** Thigmotaxis. Time spent in the margin area of the open field box was measured every 5 min for 20 min. (WT, n=10, *Auts2*$^{del8/+}$ n=10). **(D)** Spontaneous locomotor activity of mice in a novel environment was measured every 5 min for 60 min. *Auts2*$^{del8/+}$ mutant mice displayed a decrease in exploratory behavior during the first 15 min period (WT, n=19, *Auts2*$^{del8/+}$, n=16). **(E)** In open field tests, *Auts2*$^{del8/+}$ mutant mice exhibited a decrease in total distance traveled in a test field area for 5 min (right graph) whereas there was no significant difference between genotypes in time spent in an inner area (left graph) as well as the ratio of distance traveled in an inner area scored as the percentage of total distance traveled (middle graph) (WT, n=19, *Auts2*$^{del8/+}$ n=15). **(F)** Associative memory of WT and *Auts2*$^{del8/+}$ mutant mice was measured by the contextual (Context-dependent) and tone cued (Tone-dependent) fear-conditioning test 24 hrs after the conditioning phase (Conditioning). Freezing responses of *Auts2*$^{del8/+}$ mice during contextual and cued memory test were comparable to WT mice while *Auts2* mutant mice exhibited a higher response to lower nociceptive stimuli relative to WT mice (Nociceptive threshold) (WT, n=18, *Auts2*$^{del8/+}$, n=15). **(G)** Summary of the results from behavioral test battery for *Auts2*$^{neo/+}$ (Hori et al., 2015) (Hori et al., 2015) and *Auts2*$^{del8/+}$ mutant mice. Data are mean ± SEM and box-and-whisker plots (medians with interquartile range, minimum, and maximum values are represented). $^{**}P < 0.01$, $^{***}P < 0.001$, (A and B) Mann-Whitney U test, (C and D) two-way ANOVA with repeated measures, (E) unpaired t-test, (F) two-way ANOVA with repeated measures in conditioning and Mann-Whitney U test in freezing responses.

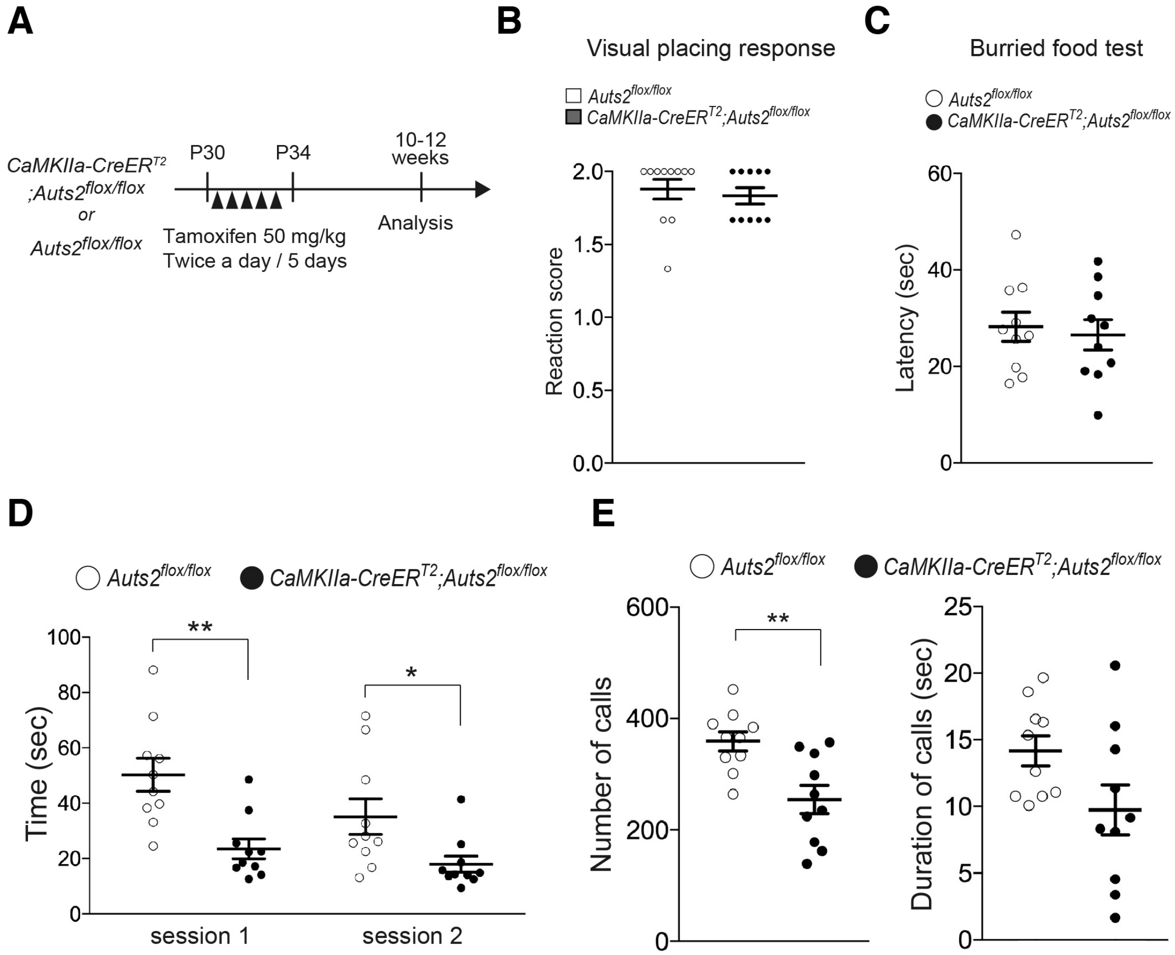
A
CaMKIIa-CreER$^{T2}$
;Auts2$^{flox/flox}$
or
Auts2$^{flox/flox}$
P30
P34
10-12 weeks
Tamoxifen 50 mg/kg
Twice a day / 5 days
Analysis
B
Visual placing response
Auts2$^{flox/flox}$
CaMKIIa-CreER$^{T2}$;Auts2$^{flox/flox}$
Reaction score
2.0
1.5
1.0
0.5
0.0
C
Burried food test
Auts2$^{flox/flox}$
CaMKIIa-CreER$^{T2}$;Auts2$^{flox/flox}$
Latency (sec)
60
40
20
0
D
Auts2$^{flox/flox}$
CaMKIIa-CreER$^{T2}$;Auts2$^{flox/flox}$
Time (sec)
100
80
60
40
20
0
**
*
session 1
session 2
E
Auts2$^{flox/flox}$
CaMKIIa-CreER$^{T2}$;Auts2$^{flox/flox}$
Number of calls
600
400
200
0
**
Duration of calls (sec)
25
20
15
10
5
0

**Supplementary Figure 5. Behavioral analyses of *CaMKIIa-CreER$^{T2}$;Auts2$^{flox/flox}$* mice. Related to Figure 6**

**(A)** Scheme illustrating the tamoxifen-inducible deletion of *Auts2* in postnatal forebrain. Tamoxifen was administered to *CaMKIIa-CreER$^{T2}$;Auts2$^{flox/flox}$* homozygotes and their control *Auts2$^{flox/flox}$* littermate mice during P30-34 and behavioral analyses were performed during 10-12 weeks. **(B)** Visual placing response test. Reaction score was rated as follow: 0, no observable placing behavior; 1, a weak or delayed placing response; 2, a clear placing reaction. (*Auts2$^{flox/flox}$*, n=11, *CaMKIIa-CreER$^{T2}$;Auts2$^{flox/flox}$* n=10). **(C)** The buried food finding test. Time spent to find buried food pellet was measured. (*Auts2$^{flox/flox}$*, n=10, *CaMKIIa-CreER$^{T2}$;Auts2$^{flox/flox}$*, n=10). **(D)** Reciprocal social interaction test. Social interaction between *Auts2$^{flox/flox}$* or *CaMKIIa-CreER$^{T2}$;Auts2$^{flox/flox}$* mouse pairs during 5 min were measured (*Auts2$^{flox/flox}$*, n=10, *CaMKIIa-CreER$^{T2}$;Auts2$^{flox/flox}$* n=10). **(E)** The number (left) and duration (right) of USVs during 1 min in adult *Auts2$^{flox/flox}$* or *CaMKIIa-CreER$^{T2}$;Auts2$^{flox/flox}$* mice (*Auts2$^{flox/flox}$*, n=10, *CaMKIIa-CreER$^{T2}$;Auts2$^{flox/flox}$* n=10). Data are the means ± SEM. *$P < 0.05$, **$P < 0.01$, ***$P < 0.001$, (B ,C and E) unpaired t-test, (D) two-way ANOVA.

Supplementary Table 1 : Summary of *Auts2* mutant Mouse Strains

| Mouse strain | Allele type | Genotype |
|---|---|---|
| *Auts2*$^{neo}$ | Global KO<br>(*Neo* gene knock-in) | WT (Control)<br>*Auts2*$^{Neo/+}$ (Heterozygote) |
| *Auts2*$^{del8}$ | Global KO<br>(Exon 8 deleted) | WT (Control)<br>*Auts2*$^{del8/+}$ (Heterozygote) |
| *Emx1*$^{Cre}$;*Auts2*$^{flox}$ | forebrain-specific conditional KO<br>(Exon 8 deleted) | *Auts2*$^{flox/flox}$ (Control)<br>*Emx1*$^{Cre/+}$;*Auts2*$^{flox/+}$ (Heterozygote)<br>*Emx1*$^{Cre/+}$;*Auts2*$^{flox/flox}$ (Homozygote) |
| *CaMKIIa-CreER*$^{T2}$;*Auts2*$^{flox}$ | Tamoxifen-inducible mature projection neuron -specific conditional KO<br>(Exon 8 deleted) | *Auts2*$^{flox/flox}$ (Control)<br>*CaMKIIa-CreER*$^{T2}$;*Auts2*$^{flox/flox}$ (Homozygote) |

### [Methods]

#### Experimental animals

*Rosa26R*$^{YFP}$ mouse line (stock no. 006148) was obtained from The Jackson Laboratory. Generation of *Auts2*$^{del8}$ and *Auts2-floxed* mice with a pure C57BL/6N genetic background and genotyping for these mice have been previously described (Supplementary Table 1) (Hori et al., 2014). *Emx1*$^{Cre}$ (stock #RBRC00808, C57BL/6J background) and *CaMKIIa-CreER*$^{T2}$ (B6.FVB-Tg(Camk2a-cre/ERT2)2Gsc/Ieg, stock #EM02125) mice were purchased from RIKEN BioResource Center (RIKEN, Tsukuba, Japan) and European Mouse Mutant Archive (EMMA) (HelmholtzZentrum München, Neuherberg, Germany), respectively (Erdmann et al., 2007; Iwasato et al., 2000). *Emx1*$^{Cre}$ mice were backcrossed for 8 generations to C57BL6/N wild type mice (Charles River Laboratories, Kanagawa, Japan) before crossing with *Auts2*$^{flox/flox}$ mice. *Emx1*$^{Cre/+}$*;Auts2*$^{flox/flox}$ homozygous mutant mice were generated by crossing *Emx1*$^{Cre/+}$ mice with *Auts2*$^{flox/flox}$ mice to yield *Emx1*$^{Cre/+}$*:Auts2*$^{flox/+}$ heterozygous mutant progeny. *Emx1*$^{Cre/+}$*;Auts2*$^{flox/+}$ male mice were then crossed with *Auts2*$^{flox/flox}$ female mice to obtain litters consisting of control (*Auts2*$^{flox/+}$or *Auts2*$^{flox/flox}$) mice, heterozygous (*Emx1*$^{Cre/+}$*;Auts2*$^{flox/+}$) or homozygous (*Emx1*$^{Cre/+}$*;Auts2*$^{flox/flox}$) mutant mice. In this study, *Auts2*$^{flox/flox}$ mice were used as the controls. For the generation of *CaMKIIa-CreER*$^{T2}$*;Auts2*$^{flox/flox}$ mice, *CaMKIIa-CreER*$^{T2}$ mice with mixed genetic background (F1: C57BL6N/FVB) were first crossed with *Auts2*$^{flox/flox}$ mice (C57BL/6N background) to obtain *CaMKIIa-CreER*$^{T2}$*;Auts2*$^{flox/+}$ mice.

*CaMKIIa-CreER$^{T2}$;Auts2$^{flox/+}$* mice were then crossed with *Auts2$^{flox/flox}$* mice to obtain litters consisting of *Auts2$^{flox/+}$*, *Auts2$^{flox/flox}$*, *CaMKIIa-CreER$^{T2}$;Auts2$^{flox/+}$* and *CaMKIIa-CreER$^{T2}$;Auts2$^{flox/flox}$* mice. For all experiments, the *CaMKIIa-CreER$^{T2}$;Auts2$^{flox/flox}$* males were crossed with *Auts2$^{flox/flox}$* females to yield the test animal cohorts consisting of the *CaMKIIa-CreER$^{T2}$;Auts2$^{flox/flox}$* and *Auts2$^{flox/flox}$* littermates. All tested animals in the behavioral analyses were generated through at least 9 crosses with C57BL/6N background animals (e.g. *Auts2$^{flox/flox}$*) to obtain genetic backgrounds close to that of C57BL/6N. For the experiments with *CaMKIIa-CreER$^{T2}$* mice, tamoxifen (Sigma-Aldrich, St. Louis, MO, USA) was administered at 50 mg/kg during postnatal 21-25 days for anatomical analysis or P30-34 for behavioral analyses by intraperitoneal injection twice daily for 5 consecutive days and the analyses were performed at postnatal day 50 and 10-12 weeks, respectively. Mice were maintained in ventilated racks under a 12-h light/dark cycle with food and water ad libitum in temperature controlled, pathogen-free facilities. Mice of each genotype were randomly allocated to different experiments.

**Behavioral analysis**

**The buried food finding test**

The buried food finding test was carried out as previously described (Yang and Crawley, 2009). Briefly, male mice were fasted for 18-24 hrs before testing. Subject mice were individually habituated in a clean cage (45 x 23 x 15 cm) for 5 min. For testing, a food pellet was buried at the end of the cage under 1 cm of wood-chip bedding. Subject mice were placed in the corner opposite to the site of the concealed food pellet. Movement of mice was recorded by video camera and time spent to explore the food pellet was measured by an examiner with stopwatch.

**Visual placing response test**

The function of the visual system was evaluated by the visual placing response as previously described by Metz and Schwab (Metz and Schwab, 2004). In this test, the test mouse was suspended by its tail and lowered toward a solid object without any contact to the vibrissae. When the head of a mouse approaches near the edge of the object, the mouse normally raises its head and extends the forelimbs to place them onto the object. The procedure was conducted by three trials and the mean response was rated with the following scoring system: 0 indicates no observable placing behavior, 1 represents a weak or delayed placing response and 2 points indicates a clear placing reaction.

**Whisker twitch reflex**

The whisker twitch reflex was tested by approaching from behind and lightly touching one set of vibrissae, eliciting head turning to the side on which the vibrissae was touched (Miyakawa et al., 2001).

**Thigmotaxis**

Mice were placed in the center of the test chamber (26 cm x 26 cm x 40 cm) under moderately bright light conditions (100 lux) and allowed to explore it. Each 20 min session was monitored by video camera and analyzed in four 5 min bins. Time spent in the marginal area defined as a 4 cm band extending from the wall was measured by examiner with a stopwatch.

**Locomotor activity**

Spontaneous exploratory locomotion was examined as described previously (Nagai et al., 2010). Mice were individually placed in a transparent acrylic cage with a black frosted Plexiglas floor (25×25×20 cm) under moderate light conditions (15 lux), and locomotor activity was measured every 5 min for 60 min using digital counters with an infrared sensor (BrainScience Idea, Osaka, Japan).

**Open field test**

Mice were placed in the center of the test chamber (diameter, 60 cm; height, 35 cm) under moderate light conditions (60 lux) and allowed to explore it for 5 min, while their activity was automatically analyzed using the ethovision automated tracking program (Brainscience Idea Co. Ltd., Osaka, Japan) (Lee et al., 2005). The center zone of the open-field was defined as the 40 cm-diameter inner circle in the chamber. Movements were measured via a camera mounted above the open field. Measurements included distance traveled and time spent in the inner and outer sections.

**Immunoblotting**

The lysates of HEK293T cells transfected with AUTS2 expression plasmids or cerebral cortices from mouse brain at P0 were solubilized in SDS sample buffer and separated in 2-15% gradient gel by SDS-PAGE (Gellex International co.ltd., Tokyo, Japan). Proteins transferred onto a PVDF membrane were immunoblotted with anti-AUTS2 antibody (HPA000390, Sigma-Aldrich, St. Louis, MO, USA) and anti-GAPDH (2118S, Cell Signaling Technology, Tokyo, Japan) antibodies, and visualized using HRP-conjugated secondary antibody (GE Healthcare, Chicago, IL, USA) followed by ECL Prime (GE Healthcare, Chicago, IL, USA). Signals were detected with a cooled CCD camera (LAS-4000 mini; Fujifilm, Kanagawa, Japan).

**Golgi-staining and analysis of dendritic spines**

Whole brains collected from mice were subjected to Golgi impregnation solution (FD Rapid GolgiStain kit, FD NeuroTechnologies, Columbia MD, USA). Coronal sections with 80-100 μm thick were obtained with cryostat and mounted on gelatin-coated slides. After tissues were processed for Golgi-Cox staining according to manufacturer's instructions, the brain sections were dehydrated with a graded series of ethanol, immersed in xylene, and embedded in Entellan (Merck, Darmstadt, Germany). Neurons were traced using a Leica microscope (DM5000B, Leica Microsystems, Danaher, Germany) with a 100x oil-immersion objective and were 3D-reconstructed by Neurolucida Software (MBF Bioscience, Williston, VT, USA). Spines along the dendritic segments (>30 μm length) within 100 μm from soma were examined, and spine densities were calculated as mean number of spines per 10 μm dendrites. On the basis of spine morphology, dendritic protrusions were classified into the following four categories (Harris and Kater, 1994): thin (≥ 0.5 μm protrusions with small bulbous head less than twice as large as spine neck), mushroom (≥0.5 μm protrusions with bulbous head more than twice as large as spine neck), stubby (≥0.5 μm protrusions with bulbous head but without a neck), filopodia (≥5 μm long and thin protrusions without bulbous heads). Dendritic protrusions with total lengths exceeding 10 μm were considered as branched dendrites and excluded from the analysis.

**Electrophysiology**

Whole-cell voltage-clamp recordings of mEPSCs and mIPSCs using brain slices were conducted as previously described (Takahashi et al., 2012). Coronal hippocampal slices with 400 μm thickness from adult mice at P33-44 were prepared in ice-cold dissection buffer (300 mM sucrose, 3.4 mM KCl, 0.6 mM $NaH_2PO_4$, 10 mM D-Glucose, 10 mM HEPES, 3.0 mM $MgCl_2$, 0.3 mM $CaCl_2$ at pH 7.4) using a VT1200S vibratome (Leica Biosystems, Danaher, Germany). Hippocampal slices were incubated in artificial cerebrospinal fluid (ACSF; 119 mM NaCl, 2.5 mM KCl, 1.0 mM $NaH_2PO_4$, 26.2 mM $NaHCO_3$, 11 mM D-Glucose, 4.0 mM $MgSO_4$, 4.0 mM $CaCl_2$, gassed with 95% $O_2$ and 5% $CO_2$), left to recover for more than 1 hour at room temperature, and then transferred to a recording chamber mounted on an upright microscope (BX61WI, Olympus, Tokyo, Japan). For voltage-clamp recordings of hippocampal slices, borosilicate glass pipettes (4-6 MΩ) were filled with the internal solutions (135 mM $CsMeSO_4$, 10 mM HEPES, 0.2 mM EGTA, 8 mM NaCl, 4 mM Mg-ATP, 0.3 mM $Na_3GTP$ at pH7.2, osmolality adjusted to 280-300 mOsm). All data of whole-cell voltage-clamp recordings were acquired with Multiclamp 700B (Molecular Devices, San Jose, CA, USA) equipped with an A/D converter (BNC-2090, National Instruments, Austin, TX, USA) and Igor Pro software version 4.01 (Wavemetrics, Portland, OR, USA) at 4 kHz. Series resistances were monitored, and the data were discarded when the series resistance changed by > 30 MΩ during recordings. mEPSCs were recorded at -70 mV in the presence of 1 μM tetrodotoxin and 100 μM picrotoxin, and mIPSCs were recorded at 0 mV in the presence of 1 μM tetrodotoxin, 10 μM CNQX and 50 μM D-APV. mEPSCs and mIPSCs events above a threshold value (10 pA) were analyzed with Minianalysis software version 6.0.3 (Synaptosoft, Fort Lee, NJ, USA).

**qPCR**

Total RNA was purified using the Qiagen RNeasy Plus Universal mini kit (QIAGEN, Hilden, Germany). Purified total RNA (0.5 μg) was subsequently reverse transcribed to cDNA using the ReverTra Ace qPCR RT kit (Toyobo, Osaka, Japan) according to the manufacturer's instructions. Real-time qPCR was performed with PowerUp SYBR Green Master Mix (Thermo Fisher Scientific, Waltham, MA, USA) in an applied Biosystems 7300 Real Time PCR System and relative expression was calculated via the 2Δ method and results were normalized to the internal control β-actin. The primers (sense and antisense, respectively) were as follows: mouse *Auts2*, 5'-AGAGCCTCTCACAGCCACTG-3' and 5'- GGTGGTGGGAGATGTGAGGA -3'; β-actin, 5'-GGCTGTATTCCCCTCCATCG-3', and 5'-CCAGTTGGTAACAATGCCATGT-3'.

**RNA-sequencing and data analysis**

Total RNA was extracted from hippocampal tissues of 4 control (*Auts2*$^{flox/flox}$) and 4 KO (*Emx1*$^{Cre/+}$;*Auts2*$^{flox/flox}$) mice at P14 using RNeasy Plus Universal Kit (QIAGEN, Hilden, Germany). Quality analyses and quantification of extracted RNA were performed using NanoDrop and Qubit Fluorometer (Thermo Fisher Scientific, Waltham, MA, USA), respectively. Sequencing libraries were prepared using the NEBNext Ultra Directional RNA Library Prep Kit for directional libraries (New England BioLabs, Tokyo, Japan) and the KAPA HTP Library Preparation Kits (KAPA Biosystems, Wilmington, MA, USA) according to the manufacturer's instructions. The RNA-seq libraries were sequenced (101 cycles) using the Illumina HiSeq platforms.

Raw sequence reads were aligned to the reference mouse genomes (GRCm38/mm10) by HISAT2 (Kim et al., 2015). Genome-wide expression levels were measured as a unit of transcripts per kilobase million (TPM) using StringTie(Pertea et al., 2015) and the numbers of reads were counted per gene per sample using htseq-count within HTSeq (Anders et al., 2015). Finally, differentially expressed genes (DEGs) were identified by DESeq2(Love et al., 2014). For gene ontology analysis, DAVID bioinformatics Resources 6.8 was used (National Institute of Allergy and Infectious Diseases-National Institute of Health; https://david.ncifcrf.gov). RNA-seq data has been deposited into GEO database with the accession number GSE134712.

### [References]


Anders, S., Pyl, P.T., and Huber, W. (2015). HTSeq--a Python framework to work with high-throughput sequencing data. Bioinformatics *31*, 166-169.

Erdmann, G., Schutz, G., and Berger, S. (2007). Inducible gene inactivation in neurons of the adult mouse forebrain. BMC Neurosci *8*, 63.

Harris, K.M., and Kater, S.B. (1994). Dendritic spines: cellular specializations imparting both stability and flexibility to synaptic function. Annu Rev Neurosci *17*, 341-371.

Hori, K., Nagai, T., Shan, W., Sakamoto, A., Abe, M., Yamazaki, M., Sakimura, K., Yamada, K., and Hoshino, M. (2015). Heterozygous Disruption of Autism susceptibility candidate 2 Causes Impaired Emotional Control and Cognitive Memory. PLoS One *10*, e0145979.

Hori, K., Nagai, T., Shan, W., Sakamoto, A., Taya, S., Hashimoto, R., Hayashi, T., Abe, M., Yamazaki, M., Nakao, K.*, et al.* (2014). Cytoskeletal regulation by AUTS2 in neuronal migration and neuritogenesis. Cell reports *9*, 2166-2179.

Iwasato, T., Datwani, A., Wolf, A.M., Nishiyama, H., Taguchi, Y., Tonegawa, S., Knopfel, T., Erzurumlu, R.S., and Itohara, S. (2000). Cortex-restricted disruption of NMDAR1 impairs neuronal patterns in the barrel cortex. Nature *406*, 726-731.

Kim, D., Langmead, B., and Salzberg, S.L. (2015). HISAT: a fast spliced aligner with low memory requirements. Nature methods *12*, 357-360.

Lee, P.R., Brady, D.L., Shapiro, R.A., Dorsa, D.M., and Koenig, J.I. (2005). Social interaction deficits caused by chronic phencyclidine administration are reversed by oxytocin. Neuropsychopharmacology *30*, 1883-1894.

Love, M.I., Huber, W., and Anders, S. (2014). Moderated estimation of fold change and dispersion for RNA-seq data with DESeq2. Genome Biol *15*, 550.

Metz, G.A., and Schwab, M.E. (2004). Behavioral characterization in a comprehensive mouse test battery reveals motor and sensory impairments in growth-associated protein-43 null mutant mice. Neuroscience *129*, 563-574.

Miyakawa, T., Yared, E., Pak, J.H., Huang, F.L., Huang, K.P., and Crawley, J.N. (2001). Neurogranin null mutant mice display performance deficits on spatial learning tasks with anxiety related components. Hippocampus *11*, 763-775.

Nagai, T., Kitahara, Y., Shiraki, A., Hikita, T., Taya, S., Kaibuchi, K., and Yamada, K. (2010). Dysfunction of dopamine release in the prefrontal cortex of dysbindin deficient sandy mice: an in vivo microdialysis study. Neurosci Lett *470*, 134-138.

Pertea, M., Pertea, G.M., Antonescu, C.M., Chang, T.C., Mendell, J.T., and Salzberg, S.L. (2015). StringTie enables improved reconstruction of a transcriptome from RNA-seq reads. Nat Biotechnol *33*, 290-295.

Takahashi, H., Katayama, K., Sohya, K., Miyamoto, H., Prasad, T., Matsumoto, Y., Ota, M., Yasuda, H., Tsumoto, T., Aruga, J., *et al.* (2012). Selective control of inhibitory synapse development by Slitrk3-PTPdelta trans-synaptic interaction. Nat Neurosci *15*, 389-398, S381-382.

Yang, M., and Crawley, J.N. (2009). Simple behavioral assessment of mouse olfaction. Curr Protoc Neurosci *Chapter 8*, Unit 8 24.
